# The Influence of Model Violation on Phylogenetic Inference: A Simulation Study

**DOI:** 10.1101/2021.09.22.461455

**Authors:** Suha Naser-Khdour, Bui Quang Minh, Robert Lanfear

## Abstract

Phylogenetic inference typically assumes that the data has evolved under Stationary, Reversible and Homogeneous (SRH) conditions. Many empirical and simulation studies have shown that assuming SRH conditions can lead to significant errors in phylogenetic inference when the data violates these assumptions. Yet, many simulation studies focused on extreme non-SRH conditions that represent worst-case scenarios and not the average empirical dataset. In this study, we simulate datasets under various degrees of non-SRH conditions using empirically derived parameters to mimic real data and examine the effects of incorrectly assuming SRH conditions on inferring phylogenies. Our results show that maximum likelihood inference is generally quite robust to a wide range of SRH model violations but is inaccurate under extreme convergent evolution.

## MainText

Markov models are commonly used in model-based phylogenetic analyses such as maximum likelihood (ML) and Bayesian inference (Felsenstein 2004; Yang 2006). A Markov model is often represented by an instantaneous rate matrix Q of size 4-by-4 for DNA or 20-by-20 for protein sequences, that describes the substitution rates between nucleotides or amino acids (henceforth denoted as states), respectively. The assumption of evolution under Markovian conditions is convenient because the probabilities of the next states only depend on the current states, independently of how the current states had evolved (Felsenstein 1981; Felsenstein 1983; Yang 1994; Swofford, et al. 1996; Yang 2006). Such approaches can be made more mathematically and computationally tractable by assuming that the Markov model is stationary, reversible, and homogeneous (SRH), assumptions which are universal to many of the most commonly-used models in phylogenetics such as those in the GTR family (Kimura 1980; Felsenstein 1981; Hasegawa, et al. 1985; Tavaré 1986; Tamura and Nei 1993; Yang 1994). Time-homogeneity means that a single Q matrix operates along all edges of the tree, i.e., all substitution rates stay constant through time. Stationarity means that the state frequencies also remain constant along all edges of the tree. Reversibility means that the substitution process remains the same in both directions.

The assumptions of homogeneity, stationarity, and reversibility imply a potential departure from biological reality (Yang and Roberts 1995; Foster and Hickey 1999; Foster 2004; Ababneh, et al. 2006). Various evolutionary processes, including mutational biases, compositional heterogeneity driven by factors such as GC-biased gene conversion, and changes in selective pressures leading to amino acid compositional shifts, on phylogenetic inference under the assumption of stationarity, reversibility, and homogeneity. Hypermutations, such as the CpG islands in mammals (Bird 1980) is an example of a non-reversible model of evolution. Reversibility assumption implies that the likelihood of a treetopology will be the same regardless of the placement of the root (Felsenstein 1981). Moreover, a reversible substitution model has up to 8 free rate parameters for nucleotides and 208 for amino acids, while a non-reversible substitution model has up to 11 free rate parameters for nucleotides and 379 for amino acids, provided that the model is still stationary (Yang 1994). These degrees of freedom can increase dramatically if the model is non-stationary or/and non-homogeneous (Jayaswal, et al. 2005; Jayaswal, et al. 2011a; Jayaswal, et al. 2011b; Jayaswal, et al. 2014) at the limit there can be an independent model of evolution on every branch of a tree, meaning that the total number of parameters is the product of the number of parameters in the substitution model and the number of branches in the tree.

Using stationary, reversible, and homogeneous substitution models to infer a phylogeny from data that has evolved under more complex conditions compromises the consistency of the ML estimation (Felsenstein 2004). Ideally, we would like to use data that comply with the assumptions of the models we apply, or alternatively, use models that are not violated by the data in hand. However, the use of non-SRH models is computationally demanding and is often not practical for large datasets. On the other hand, removing data that do not comply with the SRH assumption, as suggested by Jermiin et al. (2017), will come at the cost of losing phylogenetic information. Both studies of simulated data (Huelsenbeck and Hillis 1993; Hillis, et al. 1994; Galtier and Gouy 1998; Conant and Lewis 2001; Ho and Jermiin 2004; Jermiin, et al. 2004; Boussau and Gouy 2006) and empirical data (Galtier and Gouy 1995; Galtier and Gouy 1998; Phillips, et al. 2004; Collins, et al. 2005; Jayaswal, et al. 2005; Jayaswal, et al. 2007; Phillips, et al. 2010; Jayaswal, et al. 2011b; Nguyen, et al. 2012; Betancur-R, et al. 2013; Naser-Khdour, et al. 2019) studies have shown that applying SRH models to data that have evolved under more complex conditions may lead to significant errors in phylogenetic inference. However, most of these simulation studies have used parameters or parameter values that are not likely to reflect most empirical datasets, and sometimes represent extreme conditions such as the independent convergence of distantly-related taxa to a GC content that differs substantially from the rest of the taxa in the tree (e.g. Hillis, et al. 1994; Ho and Jermiin 2004; Jermiin, et al. 2004). While these simulations are based on biological observations such as the evolution of extreme GC content differences among closely related bacteria (Mooers and Holmes 2000), they do not represent the degree of violation of SRH conditions typical of most datasets. Indeed, apart from extreme cases (e.g. Jermiin et al. 2004) it remains relatively poorly understood to what extent different types and degrees of violations of the SRH conditions affect phylogenetic inference.

In this study, we examine the influence of violating the SRH assumptions on phylogenetic inference with SRH models using parameters and parameter values that are derived from thousands of empirical datasets. We simulate nucleotide alignments under various non-stationary (and thus non-reversible) or/and non-homogeneous conditions and examine the effects of incorrectly assuming SRH conditions on inferring phylogenies from these data. It is noteworthy that by non-homogeneous conditions we refer to heterogeneity across lineages and not heterogeneity across sites. In contrast to heterogeneity across lineages that are widely ignored in phylogenetic analyses, heterogeneity across sites is usually accounted for by using a pre-defined distribution of rate variation amongst sites, such as Gamma distribution (Yang 1994). For the purpose of this study, we assume a constant rate of substitution among sites.

## Materials andMethods

### Code and Data Availability

All scripts and code used to perform the analyses in this study are available on GitHub at: https://github.com/suhanaser/empiricalGTRdist.

Simulated sequence alignments and associated metadata are deposited in the Dryad Digital Repository, DOI: 10.5061/dryad.k3j9kd582.

### Simulations

In order to investigate the ability of SRH models to correctly infer topologies and branch lengths from non-SRH data, we devised a new approach that allows us to simulate alignments gradually ranging from true SRH conditions (with identical base frequencies and identical reversible substitution processes on every branch of the topology) to the most extreme violation with completely unrelated base frequencies and non-reversible substitution processes on every branch of the topology. For an alignment of *m* taxa and *n* sites, we will denote the set of all branches in the rooted tree τ as Φ = {1,…,*l*}.

We simulate data under two different simulation schemes as follows:

1. An inheritance scheme designed to reflect the evolutionary process, in which each node in the tree inherits its substitution processes from its parent with a constant strength of inheritance modified by the branch length connecting the two nodes. The scheme reflects the continuity of evolutionary processes that are changing through time along a phylogenetic tree. The code used to perform these simulations is available in the inheritance_Simulations.py script of our GitHub repository.
2. A two-matrix scheme designed to reflect previous approaches to simulating non-SRH evolution, where two independent subtrees (that are not sisters nor descendants of each other) have an identical substitution process that is distinct from the substitution process that operates on the rest of the tree. This scheme resembles convergent evolution. The code used to perform these simulations is available in the two_matrix_Simulations.py script of our GitHub repository.

Applying these two schemes allows us to ask how evolutionarily-inspired non-SRH simulations are affected by SRH assumptions (scheme 1) and then to directly compare these to the more extreme forms of non-SRH evolution that are more often simulated (scheme 2). We will describe both simulation approaches in more detail below. But we start by describing how we choose model parameters for our simulations.

### Estimating Empirical Parameter Distributions and Tree Topologies for Simulations

Both of our simulation approaches require us to choose base frequency vectors and rate matrices with which to simulate alignments. Generating these at random could limit the applicability of our results because it is unlikely that randomly-generated base frequency vectors or rate matrices would reflect reality. To address this, we instead estimated base frequency vectors and rate matrices from a large collection of empirical alignments and then used these parameters for our simulations.

In order to estimate the distributions of the empirical base frequencies (Π) and the substitution rates (*X*) we used 32,666 partitions from 49 nucleotide datasets (Appendix Table A.1). Consisting of different types of loci (codon positions, rR 203, Column 3NA, tRNA, introns, intergenic spacers, and UCEs) and genomes (nuclear, mitochondria, virus, plastid). Since different partitions of the genome evolve differently, for each partition, we ran IQ-TREE (Minh, et al. 2020) with a GTR model and free rate heterogeneity across sites (Yang 1995) with 4 categories + invariant sites and extracted the parameters using the script extract_GTR_parameters_dist_from_empirical_data.py script available on our GitHub repository. This gave us the distributions of 32,666 estimates of each parameter in the GTR matrix (A↔C, A↔G, A↔T, C↔G, C↔T, G↔T) and the distribution of each base frequency *(π _A_, π_C_, π_G_, π_T_*).

We use a similar approach to estimate the distribution of branch lengths. Estimating branch lengths from each partition separately could be misleading because there tends to be a high stochastic error in branch lengths estimated from short single-partition alignments (Kumar, et al. 2012). Therefore, in order to estimate the empirical distribution of the branch lengths, we instead estimated a single set of branch lengths from each of our 49 nucleotide datasets and complemented these with an additional 18 amino-acid datasets. For each dataset, we ran IQ-TREE with the best-fit fully-partitioned model (Chernomor, et al. 2016), which allows each partition to have its own evolutionary model and edge-linked rates determined by ModelFinder (Kalyaanamoorthy, et al. 2017). We then rooted the tree with the outgroup taxa (if provided) and extracted the empirical branch lengths of the ingroup (*T*) for each of the 33,178 partitions from 67 nucleotide and amino acid datasets.

Finally, for each parameter in *X* (5 parameters -G↔T equals to 1) and Π (4 parameters), and for each distribution in *T* (67 distributions - each dataset is an independent distribution) we find the best-fit distribution from 36 common probability distributions using the Kolmogorov-Smirnov test using SciPy (Virtanen, et al. 2020). Assuming that sampling among each distribution independently will approximate a sample from the joint distribution of parameters, we then sampled parameters for our simulations from these best-fit distributions. Since the parameters of Π are not independent, to sample a base-frequency vector we randomly sampled a parameter from each of the four base-frequency’s best-fit distribution and then normalized these parameters to sum to 1.

The tree topology τ is derived from birth-death simulations with speciation rate λ, extinction rate μ and the fraction of sampled taxa *f* using TreeSim package with a fixed number of extant species (Stadler 2011) using the run birth-death.R available on out GitHub repository. In principle, it is possible to estimate the speciation and extinction rates from empirical data (Nee, et al. 1994; Rannala and Yang 1996; Magallon and Sanderson 2001).However, not knowing the fraction of sampled taxa a priori will tend to bias such estimates (Stadler 2013; Hua and Lanfear 2018). Because of the challenges of reliably estimating empirical speciation and extinction rates, we instead randomly sampled the speciation rate, the extinction rate and the fraction of sampled taxa from uniform distributions, to attempt to cover all the realistic regions of the parameter space.

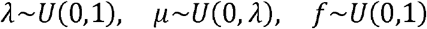

Note that under these conditions λ is always greater than μ.

We simulated datasets with 20, 40, 60, 80 and 100 taxa. For each number of taxa (*m*), we simulated 3960 topologies with random speciation rate (λ), random extinction rate (μ) and random fraction of sampled taxa (*f*). For each of these topologies, we then randomly choose a distribution from set *T* and sampled the branch lengths from this distribution (2*m*–2 branch lengths in total).

Other Python libraries that we used for the simulations are NumPy (Walt, et al. 2011), pandas (McKinney 2010) and ETE3 (Huerta-Cepas, et al. 2016). The python scripts for all simulations can be found on Github (https://github.com/suhanaser/empiricalGTRdist).

## Inheritance Evolution: Inheritance Scheme Simulations

An evolutionary scenario would, ideally, have each lineage inheriting the parameters of its molecular evolutionary process from its parent lineage. At one extreme – where inheritance is perfect and the original evolutionary process is SRH, such a process would define a molecular evolutionary process that is SRH across the entire topology by simply defining a single SRH model at the root node. At the other extreme, where the association between parent and offspring lineages is no better than random and the original process is not SRH, there is no association between parent and offspring lineages and the process is maximally non-SRH. To mimic this situation, we designed a simulation approach that allows us to vary the homogeneity and stationarity assumptions both independently and together.

Our inheritance scheme allows us to vary the degree to which a single alignment has evolved under SRH conditions by simply adjusting the strength of inheritance of the substitution process and the base frequencies either jointly via a parameter we call *ρ*, or independently via parameters ω and *ν* respectively. When the inheritance parameters are set to 1 and the model at the root of the tree is reversible, the model will conform to SRH conditions. We can simulate increasing violation of SRH conditions simply by decreasing the inheritance parameters towards zero. When the relevant inheritance parameter is less than one, each branch inherits some proportion of its substitution model from the parent branch, while the remaining proportion of the model is selected at random from the empirical parameter distributions. In practice, the parameter in a descendant branch is calculated as the weighted sum of the parameter in the parent branch (where the weight is the inheritance parameter) and a randomly-generated parameter from the appropriate empirical distribution (where the weight is one minus the inheritance parameter).

We simulated data under five different categories of conditions using this scheme, in order to examine independently and together the effects of relaxing the stationarity and homogeneity assumptions.

### 1. SRH conditions (Fig. 1a)

In the simplest case for a model that conforms to the SRH assumptions, where model parameters are generated from the empirical distributions. This describes a model in which all branches inherit this reversible model from their parent branch without variation, such that all branches on the tree have the same reversible substitution model, conforming to the SRH assumptions.

**Figure 1.**
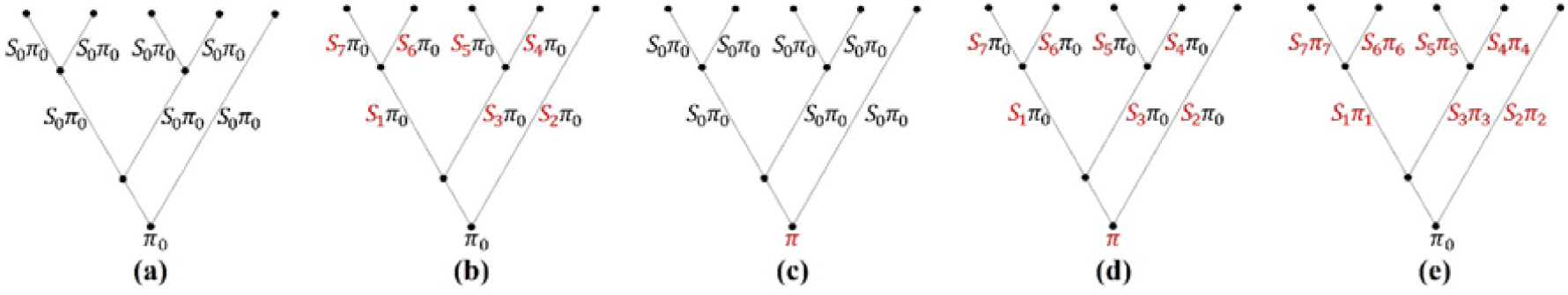
An example of 5 taxon tree with different degrees for homogeneity and stationarity. **(a)** stationary and homogeneous, **(b)** stationarity but not homogeneous, **(c)** non-stationary but homogeneous, **(d)** non-stationary and non-homogeneous where the stationarity and homogeneity assumptions are relaxed simultaneously but independently, **(e)** non-stationary and non-homogeneous where the stationarity and homogeneity assumptions are relaxed jointly.

### 2. Relaxing the stationarity assumption (Fig. 1b)

In order to hold the homogeneity assumption but relax the stationarity assumption, we introduce a parameter called ν (0 ≤ ν ≤ 1) that allows the state frequency to vary at the root while still keeping the same rate matrix for all branches of the tree. Mathematically, this can be described as:

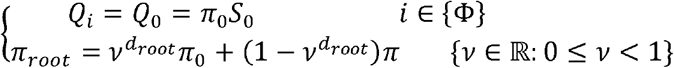

Where *Q_i_* is the substitution rate matrix operating on branch *i*, π_0_ is the stationary base frequencies, *π_root_* is the base frequencies at the root, and *d_root_* is the branch length of the root branch.

When *ν* = 1, *π_root_* is equal to *π*_0_ and this scheme boils down to the first SRH condition. When *ν* = 0, *π_root_* is equal to *π*, meaning that the root frequency is generated separately from *π*_0_. *π_root_* will vary between these two extremes when *ν*is between 0 and 1, with lower *ν* reflecting a larger deviation from stationary conditions.

### 3. Relaxing the homogeneity assumption (Fig. 1c)

In order to hold the stationarity assumption but relax the homogeneity assumption we need to simulate data in which *ν* is set to 1 (such that all branches have the same base frequencies as the root node), but we introduce a parameter ω that varies between zero and one (such that the inheritance of the parameters of the *Q* matrix ranges from completely random to near-perfect). We can describe this mathematically as follows:

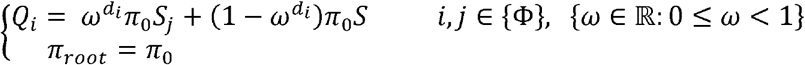

Where *Q_i_* is the process operating on branch i,*S_j_*are the substitution rates on the parent branch of branch *i*, and *d_i_* is the branch length of the branch *i*.

### 4. Relaxing the stationarity and homogeneity assumptions simultaneously but independently (Fig. 1d)

We can simulate non-stationary and non-homogeneous data by setting both *ν* and ω to values less than one. When we relax both assumptions, we will allow *Q_i_* and *π_root_* to vary simultaneously but independently:

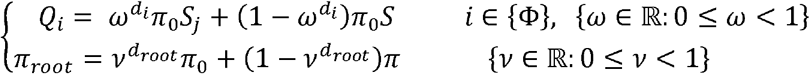

### 5. Relaxing the stationarity and homogeneity assumptions jointly (Fig. 1e)

While the 4^th^ set of simulation conditions, above, allows us to vary homogeneity and stationarity jointly but independently, it suffers from the limitation that we have a maximum of two base frequency vectors in the tree (*π_root_* and *π_0_*). To relax this assumption further, we will allow *Q_i_* to vary while *π_root_* stays fixed. In those settings, both homogeneity and stationarity will increase with *ρ*.

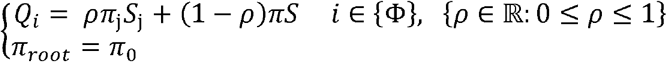

#### Convergent Evolution: The Two-Matrix Scheme Simulations

Previous studies for simulating non-SRH evolution on phylogenies have used an approach in which two distantly related branches undergo severe but correlated changes in the molecular evolutionary process. To compare this approach to the more evolutionarily-motivated approach described above, we randomly chose two nodes that are not sisters and not descendants of each other and assigned a different rate matrix (denoted by *S*_1_ *π*_1_) from the rest of the tree to all their descendant branches (Fig. 2Figure2).

**Figure 2.**
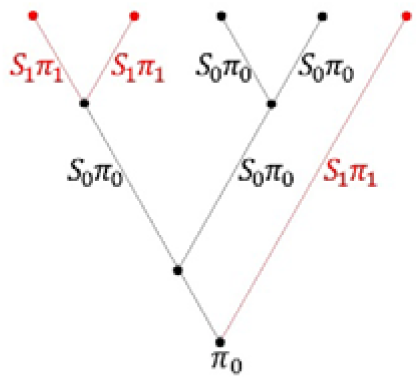
Non-stationary and non-homogeneous process with two different rate matrices Q_0_ and Q_1_.

#### Simulation Parameters

The simulation parameters that we use in this study are the strength of inheritance of the substitution process *(ω*), strength of inheritance of the base frequencies (**ν**), strength of inheritance of the substitution process and base frequencies (*ρ*), number of sites (*n*), and number of taxa (*m*) where the parameter space is:

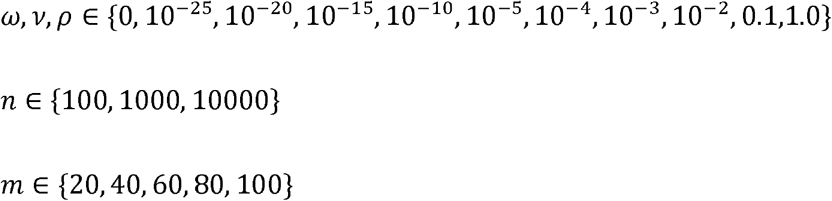

The inheritance weight parameters (ω,**ν*,ρ*) were chosen to represent an even spread of corrected inheritance weights (i.e., the inheritance weights raised to the power of *d*, where *d* is the branch length) between zero and one. The number of taxa and number of sites are chosen to reflect typical sizes of empirical datasets. For simulation under the inheritance scheme, we simulated 10 alignments of each combination of *n,m, *ν**, and ω or *n,m*, and *ρ* for a total of 19,800 simulations. For simulation under the two-matrix scheme, we simulated 1000 alignments of each combination of *n* and *m* for a total of 15,000 simulations.

#### Tree Inference

Our first goal is to understand how the incorrect use of SRH models on data that have evolved under non-SRH processes can affect phylogenetic inference. To do this, we compare the tree topologies and branch lengths estimated with SRH models in IQ-TREE to the topologies and branch lengths used to simulate each dataset. For each simulated alignment, we ran IQ-TREE with ModelFinder (Kalyaanamoorthy, et al. 2017) and 1000 ultrafast bootstrap replicates (Hoang, et al. 2018). In order to assess the ability of SRH models to infer the correct tree topology we then compared the simulated tree topology to the estimated tree topology using three different metrics – normalized Robinson-Foulds distance (Robinson and Foulds 1981), Quartet distance (Estabrook, et al. 1985), and the Path-difference distance (Steel and Penny 1993). The normalized Robinson-Foulds distance between two trees is the fraction of internal branches that appear in one tree but not the other. It ranges from 0 to 1, where 0 means that the two trees are topologically identical and 1 means that the two trees have no splits in common. In order to assess the accuracy of branch length estimates, we tested whether the estimated branch lengths and the original branch lengths are drawn from the same distribution using the two-sample Kolmogorov-Smirnov test.

## Results

### Empirical Distributions

We derived the empirical distributions of the substitution model parameters, the nucleotide frequencies, and the proportion of invariant sites from 32,666 nucleotide alignments (Appendix Table A.2). The empirical distribution of branch lengths we derived from 67 nucleotide and amino acid alignments consisting 33,178 partitions (Appendix Table A.1).

Using Kolmogorov-Smirnov test, we found the best-fit probability distribution for each one of these empirical distributions (Table 1, Appendix Table A.2, Figs. A.1-3).

**Table 1.**
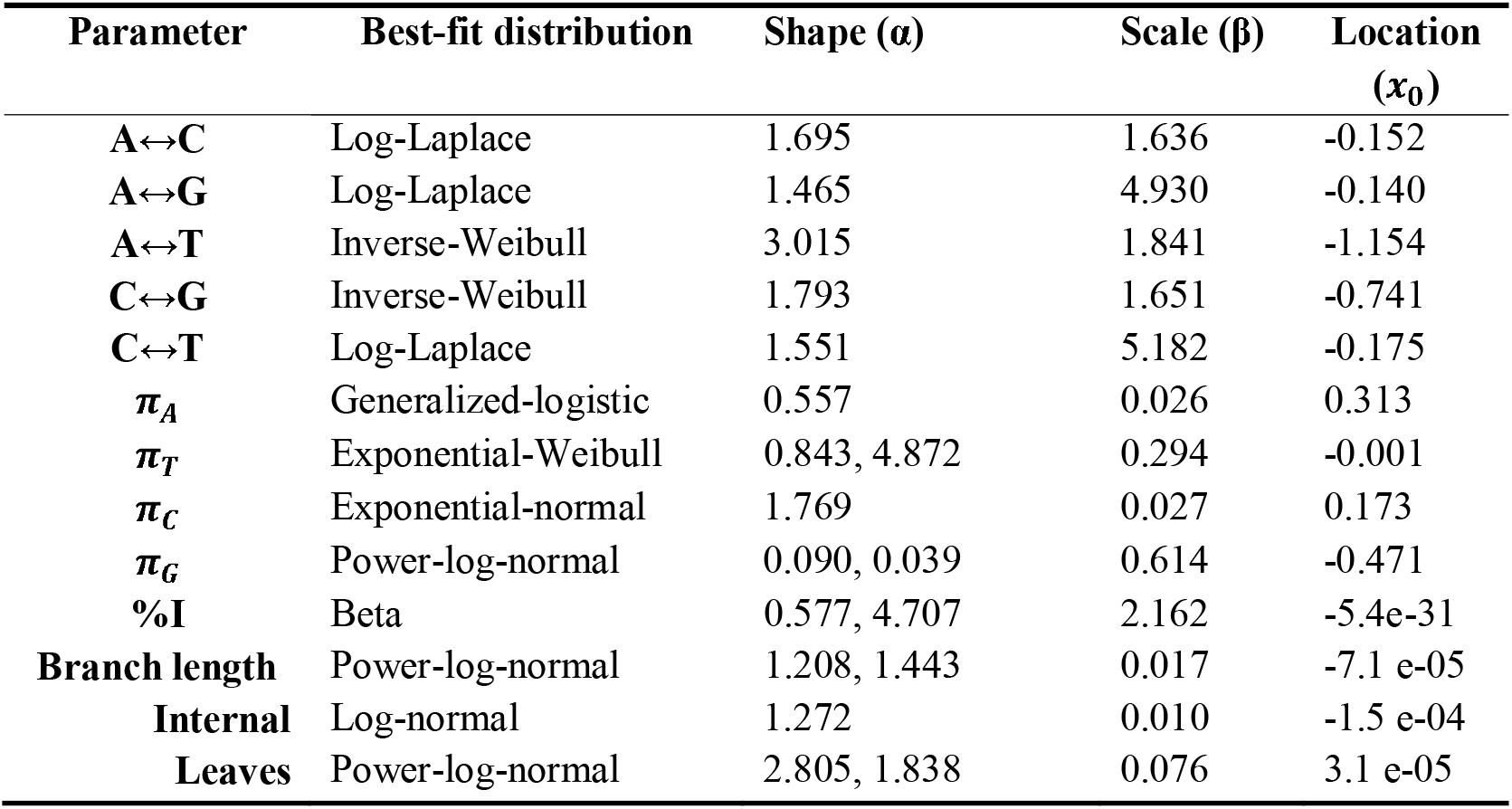
The best-fit probability distribution by Kolmogorov-Smirnov test.

| Parameter | Best-fit distribution | Shape ($\alpha$) | Scale ($\beta$) | Location ($x_0$) |
| --- | --- | --- | --- | --- |
| A↔C | Log-Laplace | 1.695 | 1.636 | -0.152 |
| A↔G | Log-Laplace | 1.465 | 4.930 | -0.140 |
| A↔T | Inverse-Weibull | 3.015 | 1.841 | -1.154 |
| C↔G | Inverse-Weibull | 1.793 | 1.651 | -0.741 |
| C↔T | Log-Laplace | 1.551 | 5.182 | -0.175 |
| $\pi_A$ | Generalized-logistic | 0.557 | 0.026 | 0.313 |
| $\pi_T$ | Exponential-Weibull | 0.843, 4.872 | 0.294 | -0.001 |
| $\pi_C$ | Exponential-normal | 1.769 | 0.027 | 0.173 |
| $\pi_G$ | Power-log-normal | 0.090, 0.039 | 0.614 | -0.471 |
| %I | Beta | 0.577, 4.707 | 2.162 | -5.4e-31 |
| **Branch length** | Power-log-normal | 1.208, 1.443 | 0.017 | -7.1 e-05 |
| **Internal** | Log-normal | 1.272 | 0.010 | -1.5 e-04 |
| **Leaves** | Power-log-normal | 2.805, 1.838 | 0.076 | 3.1 e-05 |

Furthermore, we checked if the distributions of the internal branch lengths and the leaf branch lengths are similar. Our results show the distribution of the leaf branch lengths is similar to the general distribution of all branches but with different parameters. On the other hand, the distribution of the internal branch lengths is different (Appendix Fig. A.3). Yet, for the purpose of this study, we will use the general branch length distribution for all branches.

### Both Simulation Schemes Capture the True extent of Compositional Heterogeneity

In order to confirm that our simulation settings are producing compositional variance patterns that are of the same order of magnitude as what we observed in the empirical data, we compared the variance in GC content between the simulated datasets and the empirical datasets.

Our results confirm that the compositional variance in both simulation schemes is indeed similar to the empirical data (Fig. 3).

**Figure 3.**
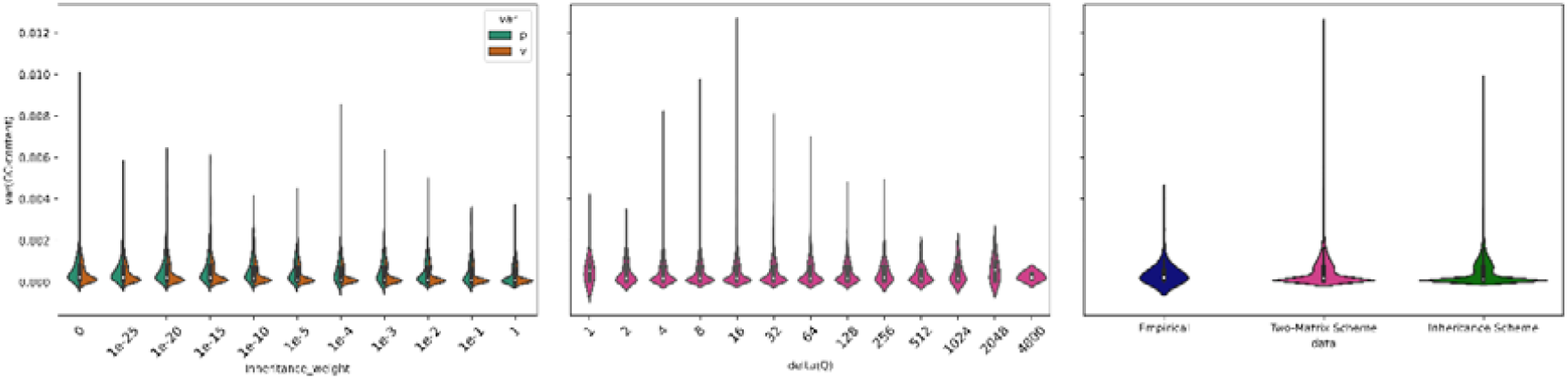
Variance in GC content in simulated and empirical datasets. The variance in GC content was calculated between sequences within each dataset.

### Phylogenetic Inference is Unaffected by Violation of SRH Conditions in an Inheritance Framework

Surprisingly, our results for the inheritance simulation scheme show that there is no detectable relationship between the severity with which SRH conditions were violated during the simulations and the accuracy of the tree topology or the tree length inferred from the simulated data. Specifically, we saw no relationship between the inheritance weight and the normalized RF (Robinson-Foulds), QD (Quartet Distance), or NPD (Normalized Path Difference) metrics in any of our inheritance simulations (Fig. 4, Appendix Figs. A.4-7). These metrics measure the difference between the inferred tree and the tree from which the alignment was simulated. If a stronger violation of the SRH conditions affects phylogenetic inference we should expect to see that the distances are higher when the inheritance weight is lower, because a lower inheritance weight implies a stronger model violation through less homogeneity (for the rate matrix) and less stationarity (for the base frequencies). It is important to note that the results presented in Fig. 4 represent averages across datasets with varying numbers of taxa and sites. While Fig. A.5 represents the non-normalized RF metric, since the sub-figures all have the same number of taxa, nRF and RF are essentially the same.

**Figure 4.**
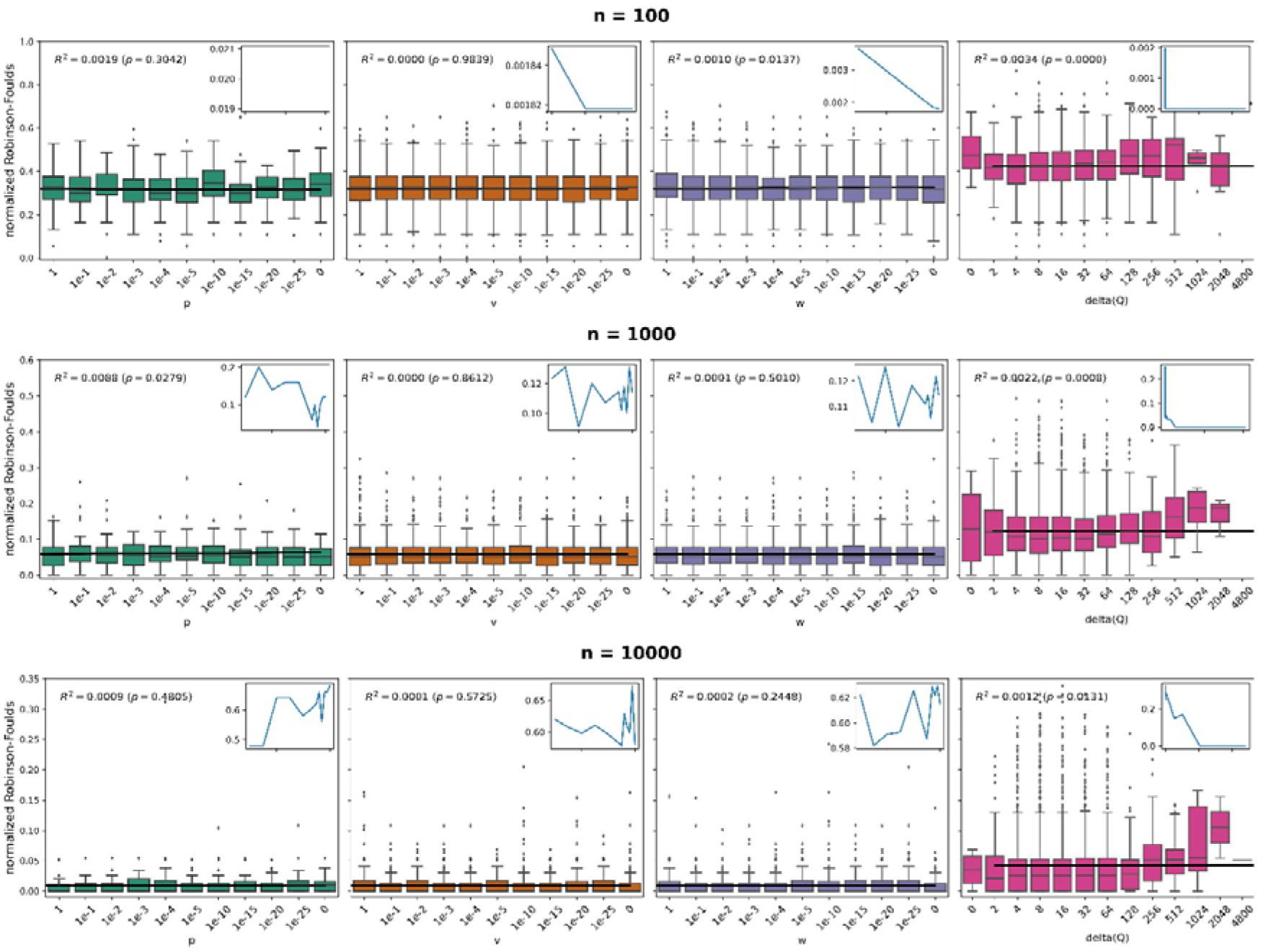
Normalized Robinson-Foulds distance between the estimated tree topology and the original tree topology as a function of the inheritance weight (*ν*, ω,ρ) in the first simulation scheme, and the distance between the two matrices (delta(Q)) in the second simulation scheme, for different number of sites. The black line is the relevant linear regression. The small plots show the proportion of datasets in which the distance between the estimated topology and the original topology equals zero as a function of the inheritance weight and the distance between the two matrices. of each inset plot matches that of the bigger plot, labels have been excluded for clarity. If violation of SRH model assumptions increases topological error, we expect the nRF distance to increase towards the right of each plot. The figure shows that for the first simulation scheme, which mimics an inheritance evolutionary process, there is no detectable association between violation of SRH conditions and topological error. For the second simulation scheme, which mimics an extreme convergent situation, topological error increases with increasing violation of SRH conditions.

In addition, our results show that the proportion of simulated datasets for which the simulated tree is recovered from the simulated alignment is constant at around 0.25 in the inheritance scheme simulations regardless of the inheritance weight (Fig. 4, Appendix Fig. A.4). Finally, we see no correlation between the inheritance weights and having a significant p-value in a Kolmogorov-Smirnov test comparing the true and estimated branch lengths (slope = 0.52, R^2^ = 0.06, p-value = 0.22), suggesting that violation of SRH assumptions in our evolutionary framework has no detectable effect on the estimation of branch lengths (Fig. 5, Appendix Fig. A.11).

**Figure 5.**
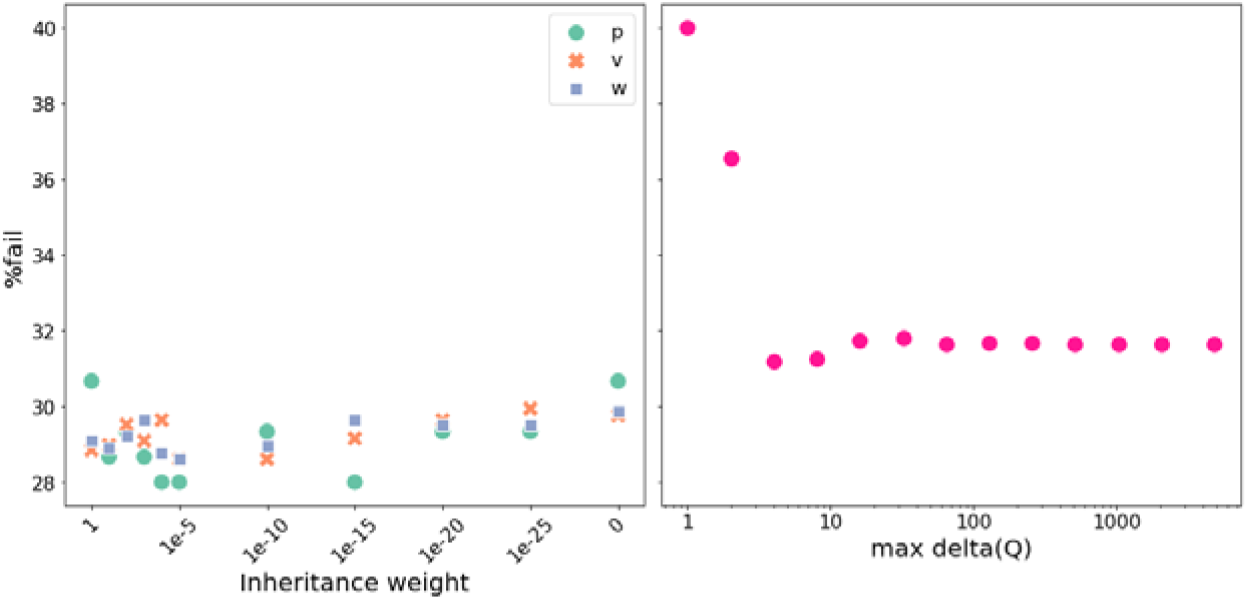
The percentage of datasets that pass the KS test (a dataset passes when there is no evidence to suggest that the inferred branch length distributions differ) as a function of the inheritance weights (*ν*,ρ,ω) (left-hand panel) and maximum Euclidean distance between the two matrices used to simulate the data (right-hand panel).

### Tree Topologies, but not Branch Lengths, are Affected by Severe and Convergent Violation of SRH Conditions

Our results show that convergent violation of SRH assumptions by allowing two distantly related branches to have identical substitution models has increasingly severe effects on phylogenetic inference as the severity of the changes in the substitution models increases. Under the two-matrix scheme, we expect to see higher distances between the true tree and the estimated tree when there are larger Euclidean distances between the original matrix and the matrix under which the divergent clades evolve. In two out of the three metrics (Robison-Foulds and Path-Difference) we found a weak but significant correlation between the distance between the matrices and the distance between the topologies (Fig. 4, Appendix Fig. A.4, Appendix Figs. A.8-10). However, in the third metric (Quartet Distance) we found no correlation. Notably, the distance between the true tree and the estimated tree increases only when the Euclidean distance between the two matrices is very high. Nevertheless, the proportion of simulated datasets for which the simulated tree is recovered from the simulated alignment declines exponentially as the difference between the matrices in the two-matrix scheme increases (Fig. 4).

The proportion of analyses in which the simulated tree is recovered positively declines from around 0.20 when there is no model violation to zero when the Euclidean distance between the matrices is around 2000, confirming that even the lowest levels of SRH violation have detectable negative effects on phylogenetic inference under the two-matrix scheme.

Finally, we see no correlation between the Euclidean distance between the two matrices and failing or passing the Kolmogorov-Smirnov test comparing the distributions of the true and estimated branch lengths (slope = 3.8E-4, R^2^ = 0.04, p-value = 0.51), suggesting that violation of SRH assumptions in the convergent framework has limited effects on the estimation of branch lengths (Fig. 5, Appendix Fig. A.12, Supplementary material tables A.2).

## Discussion

Using two different simulation schemes, we explored the impact of violating the assumption of evolution under stationary, reversible, and homogeneous (SRH) conditions on ML phylogenetic tree inference. Our study extends the simulations in many previous studies by simulating data under an evolutionary scenario in which molecular evolutionary models evolve along a phylogeny. Our results show that the inference of phylogenetic tree topologies and branch lengths are surprisingly robust to violations of SRH assumptions under an evolutionary scheme. But similarly to previous studies, we show that in extreme cases of convergent molecular evolution, the incorrect assumption of SRH conditions can severely mislead phylogenetic inference.

The first simulation scheme we introduced in this paper, which we called *the inheritance scheme*, allows tree branches to inherit their substitution process from their ancestor. The second simulation scheme, which we called *the two-matrix scheme*, is similar to previous studies and allows two distantly related monophyletic sub-trees to evolve with a different evolutionary process from the rest of the tree (Galtier and Gouy 1995; Jermiin, et al. 2004; Jayaswal, et al. 2011a; Duchene, et al. 2017).

Surprisingly, our results show no correlation between errors in the topology or branch length distribution inference and any of the inheritance scheme parameters, even in extreme cases where the evolutionary process is completely heterogeneous and non-stationary. These results indicate that ML tree inference with SRH models is surprisingly robust to even quite extreme violations of the SRH conditions.

Under the two-matrix simulation scheme, we found a small but significant increase in topological inference error and the extent of the violation of the SRH assumptions. Specifically, the more extreme the evolutionary convergence, the larger the errors in topological inference that assume SRH conditions. Despite this, we found no correlation between branch length inference and the distance between the two matrices. These results emphasize the limitations of ML inference to operate under certain model violations, especially when these violations are highly imbalanced along the tree, as in the case of the two-matrix scheme simulations. These results indicate that the inference of the substitution model is more influenced by the imbalance of the model violation distribution along the tree than by the model violation itself. This conclusion agrees well with all previous simulation studies of similar simulation conditions (e.g. Conant and Lewis 2001; Jermiin, et al. 2004; Duchene, et al. 2017).

It is noteworthy that our results from the different simulation schemes agree with the results from empirical data (Naser-Khdour, et al. 2019). When the model violations were more severe, like in the two-matrix scheme, we obtained a significant correlation between topological inference error and the extent of the violation of the SRH assumptions. In the inheritance scheme simulations, on the other hand, the violation of the SRH assumptions was milder, and therefore the phylogenetic inference was more robust. These results indicate that the extent of the violation of the SRH assumptions in the empirical datasets was higher than in our simulations under the inheritance scheme and it was probably closer to the extent of violation in the two-matrix scheme simulations. Moreover, they emphasize the impact of model violation due to non-SRH evolution on phylogenetic inference and suggest reducing model violation in phylogenetic analysis by using the new protocol of phylogenetic inference (Jermiin, et al. 2020) or using more complex substitution models e.g.(Galtier and Gouy 1998; Tamura and Kumar 2002; Blanquart and Lartillot 2008; Dutheil, et al. 2012; Zou, et al. 2012; Groussin, et al. 2013; Jayaswal, et al. 2014) has the potential to improve phylogenetic accuracy.

For the purpose of this study, in order to simulate data that mimic as closely as possible empirical alignments, we extracted the empirical distributions of base frequencies, substitution rates, the proportion of invariable sites, and branch lengths from tens of thousands of empirical datasets. In addition to their use in this paper, these empirical distributions, along with their best-fit distributions may be useful for a wide variety of simulation studies, or for specifying prior distributions for Bayesian phylogenetic methods.

