## Appendix for "The Influence of Model Violation on Phylogenetic Inference: A Simulation Study"

### Tables

#### Table A.1. Number of Taxa, Number of Sites, Type, Clade, and Study Reference for Each Data Set That Have Been Used in This Study

|  | **Dataset** | **Study Reference** | **Dataset Reference** | **Type** | **Clade** | **Taxa** | **Sites** | **Partitions** |
| --- | --- | --- | --- | --- | --- | --- | --- | --- |
| 1 | Anderson_2013 | (Anderson, et al. 2014) | (Anderson, et al. 2013) | DNA | Loliginidae | 145 | 3037 | 4 |
| 2 | Ballesteros_2019 | (Ballesteros and Sharma 2019a) | (Ballesteros and Sharma 2019b) | AA | Chelicerata | 53 | 1484206 | 3534 |
| 3 | Becker_2016 | (Becker, et al. 2016) | (Becker, et al. 2017) | AA | Halobacteriacea | 170 | 217 | 1 |
| 4 | Bergsten _2013 | (Bergsten, et al. 2013a) | (Bergsten, et al. 2013b) | DNA | Dytiscidae | 38 | 2111 | 8 |
| 5 | Borowiec_2015 | (Borowiec, et al. 2015) | (Borowiec, et al. 2016) | AA | Metazoa | 36 | 384981 | 1080 |
| 6 | Branstetter_2017 | (Branstetter, et al. 2017b) | (Branstetter, et al. 2017a) | DNA | Aculeata | 187 | 183747 | 807 |
| 7 | Broughton_2013 | (Broughton, et al. 2013b) | (Broughton, et al. 2013a) | DNA | Osteichthyes | 61 | 19997 | 61 |
| 8 | Brown_2012 | (Brown, et al. 2012b) | (Brown, et al. 2012a) | DNA | Ptychozoon | 41 | 1665 | 7 |
| 9 | Cannon_2016a | (Cannon, et al. 2016a) | (Cannon, et al. 2016b) | AA | Metazoa | 78 | 44896 | 212 |
| 10 | Cannon_2016b | (Cannon, et al. 2016a) | (Cannon, et al. 2016b) | DNA | Metazoa | 78 | 89792 | 424 |
| 11 | Chen_2015 | (Chen, et al. 2015b) | (Chen, et al. 2015a) | AA | Gnathostomata | 58 | 1806035 | 4682 |
| 12 | Cognato_2001 | (Cognato and Vogler 2001b) | (Cognato and Vogler 2001a) | DNA | Coleoptera: Scolytinae | 44 | 1897 | 7 |
| 13 | Crawford_2012 | (Crawford, et al. 2012b) | (Crawford, et al. 2012a) | DNA | Sauria | 10 | 465241 | 1145 |
| 14 | Day_2013 | Day, et al. (2013a) | (Day, et al. 2013b) | DNA | Synodontis | 152 | 3586 | 11 |
| 15 | Devitt_2013 | (Devitt, Devitt, et al. 2013) | (Devitt, Cameron Devitt, et al. 2013) | DNA | Ensatina eschscholtzii klauberi | 69 | 823 | 4 |
| 16 | Dornburg_2012 | (Dornburg, et al. 2012b) | (Dornburg, et al. 2012a) | DNA | Teleostei: Beryciformes: Holocentridae | 44 | 5919 | 21 |
| 17 | Faircloth_2013 | (Faircloth, et al. 2013b) | (Faircloth, et al. 2013a) | DNA | Actinopterygii | 27 | 149366 | 491 |
| 18 | Fong_2012 | (Fong, et al. 2012b) | (Fong, et al. 2012a) | DNA | Vertebrata | 110 | 25919 | 168 |
| 19 | Horn_2014 | (Horn, et al. 2014b) | (Horn, et al. 2014a) | DNA | Euphorbia | 197 | 11587 | 28 |
| 20 | Irisarri_2017 | (Irisarri, et al. 2017b) | (Irisarri, et al. 2017a) | AA | Gnathostomata | 100 | 1964439 | 4593 |
| 21 | Jarvis_2015 | (Jarvis, et al. 2015) | (Jarvis, et al. 2014) | AA | Aves | 52 | 4519041 | 8295 |
| 22 | Kawahara_2013 | (Kawahara and Rubinoff 2013a) | (Kawahara and Rubinoff 2013b) | DNA | Hyposmocoma | 70 | 2238 | 9 |
| 23 | Lartillot_2012 | (Lartillot and Delsuc 2012b) | (Lartillot and Delsuc 2012a) | DNA | Eutheria | 78 | 15117 | 51 |
| 24 | Leache_2015 | (Leache, et al. 2015) | (Leaché, et al. 2015) | DNA | Phrynosomatinae | 11 | 358363 | 583 |
| 25 | Looney_2016 | (Looney, et al. 2016) | (Looney, et al. 2015) | DNA | Russula | 1171 | 3927 | 4 |
| 26 | McCormack_2013 | (McCormack, et al. 2013b) | (McCormack, et al. 2013a) | DNA | Neoaves | 33 | 539526 | 1541 |
| 27 | Meiklejohn_2016 | (Meiklejohn, et al. 2016a) | (Meiklejohn, et al. 2016b) | DNA | Phasianidae | 18 | 614159 | 1501 |
| 28 | Misof_2014 | (Misof, et al. 2014b) | (Misof, et al. 2014a) | AA | Insecta | 144 | 595033 | 2868 |
| 29 | Moyle_2016 | (Moyle, et al. 2016b) | (Moyle, et al. 2016a) | DNA | Oscines | 106 | 375172 | 515 |
| 30 | Murray_2013 | (Murray, et al. 2013a) | (Murray, et al. 2013b) | DNA | Eucharitidae | 237 | 3111 | 9 |
| 31 | Near_2013 | (Near, et al. 2013b) | (Near, et al. 2013a) | DNA | Acanthomorpha | 608 | 8577 | 30 |
| 32 | Nguyen_2016a | (Nguyen, et al. 2016b) | (Nguyen, et al. 2016a) | AA | Hymenoptera | 17 | 688 | 1 |
| 33 | Nguyen_2016b | (Nguyen, et al. 2016b) | (Nguyen, et al. 2016a) | AA | Hymenoptera | 31 | 680 | 1 |
| 34 | Nguyen_2016c | (Nguyen, et al. 2016b) | (Nguyen, et al. 2016a) | AA | Hymenoptera | 25 | 811 | 1 |
| 35 | Nguyen_2016d | (Nguyen, et al. 2016b) | (Nguyen, et al. 2016a) | AA | Hymenoptera | 17 | 704 | 1 |
| 36 | Nguyen_2016e | (Nguyen, et al. 2016b) | (Nguyen, et al. 2016a) | AA | Hymenoptera | 17 | 385 | 1 |
| 37 | Nguyen_2016f | (Nguyen, et al. 2016b) | (Nguyen, et al. 2016a) | AA | Hymenoptera | 17 | 583 | 1 |
| 38 | Oaks_2011 | (Oaks 2011b) | (Oaks 2011a) | DNA | Crocodylia | 79 | 7282 | 50 |
| 39 | Prebus_2017 | (Prebus 2017b) | (Prebus 2017a) | DNA | Temnothorax | 50 | 1561581 | 2098 |
| 40 | Pyron_2011 | (Pyron and Wiens 2011) | (Pyron, et al. 2011) | DNA | Amphibia | 2872 | 12712 | 34 |
| 41 | Ran_2018a | (Ran, et al. 2018b) | (Ran, et al. 2018a) | AA | Spermatophyta | 38 | 432014 | 1308 |
| 42 | Ran_2018b | (Ran, et al. 2018b) | (Ran, et al. 2018a) | DNA | Spermatophyta | 38 | 1296042 | 3924 |
| 43 | Reddy_2017 | (Reddy, et al. 2017b) | (Reddy, et al. 2017a) | DNA | Aves | 235 | 137324 | 88 |
| 44 | Richart_2015 | (Richart, et al. 2016b) | (Richart, et al. 2016a) | DNA | Ischyropsalidoidea | 6 | 536124 | 2016 |
| 45 | Rightmyer_2013 | (Rightmyer, et al. 2013b) | (Rightmyer, et al. 2013a) | DNA | Hymenoptera: Megachilidae | 94 | 3692 | 25 |
| 46 | Sauquet_2011 | (Sauquet, et al. 2012) | (Sauquet, et al. 2011) | DNA | Nothofagus | 51 | 5444 | 10 |
| 47 | Seago_2011 | (Seago, et al. 2011b) | (Seago, et al. 2011a) | DNA | Coccinellidae | 97 | 2253 | 7 |
| 48 | Sharanowski_2011 | (Sharanowski, et al. 2011b) | (Sharanowski, et al. 2011a) | DNA | Braconidae | 139 | 3982 | 11 |
| 49 | Shen_2018 | (Shen, et al. 2018) | (Shen 2018) | AA | Saccharomycotina | 343 | 1162805 | 2407 |
| 50 | Siler_2013 | (Siler, Oliveros, et al. 2013) | (Siler, Brown, et al. 2013) | DNA | Lycodon | 61 | 2697 | 7 |
| 51 | Smith_2014 | (Smith, et al. 2014b) | (Smith, et al. 2014a) | DNA | Xenops minutus | 8 | 825804 | 1366 |
| 52 | Tolley_2013 | (Tolley, et al. 2013b) | (Tolley, et al. 2013a) | DNA | Chamaeleonidae | 203 | 5054 | 16 |
| 53 | Unmack_2013 | (Unmack, et al. 2013b) | (Unmack, et al. 2013a) | DNA | Melanotaeniidae | 139 | 6827 | 25 |
| 54 | Varga_2019 | (Varga, et al. 2019b) | (Varga, et al. 2019a) | DNA | Basidiomycota | 5285 | 5737 | 3 |
| 55 | Wainwright_2012 | (Wainwright, et al. 2012a) | (Wainwright, et al. 2012b) | DNA | Acanthomorpha | 188 | 8439 | 30 |
| 56 | Whelan_2017 | (Whelan, et al. 2017a) | (Whelan, et al. 2017b) | AA | Metazoa | 76 | 49388 | 127 |
| 57 | Wood_2012 | (Wood, et al. 2013) | (Wood, et al. 2012) | DNA | Archaeidae | 37 | 5185 | 8 |
| 58 | Worobey_2014a | (Worobey, et al. 2014b) | (Worobey, et al. 2014a) | DNA | Influenzavirus A | 146 | 1716 | 3 |
| 59 | Worobey_2014b | (Worobey, et al. 2014b) | (Worobey, et al. 2014a) | DNA | Influenzavirus A | 327 | 759 | 3 |
| 60 | Worobey_2014c | (Worobey, et al. 2014b) | (Worobey, et al. 2014a) | DNA | Influenzavirus A | 92 | 1416 | 3 |
| 61 | Worobey_2014d | (Worobey, et al. 2014b) | (Worobey, et al. 2014a) | DNA | Influenzavirus A | 355 | 1497 | 3 |
| 62 | Worobey_2014e | (Worobey, et al. 2014b) | (Worobey, et al. 2014a) | DNA | Influenzavirus A | 340 | 699 | 3 |
| 63 | Worobey_2014f | (Worobey, et al. 2014b) | (Worobey, et al. 2014a) | DNA | Influenzavirus A | 332 | 2151 | 3 |
| 64 | Worobey_2014g | (Worobey, et al. 2014b) | (Worobey, et al. 2014a) | DNA | Influenzavirus A | 326 | 2274 | 3 |
| 65 | Worobey_2014h | (Worobey, et al. 2014b) | (Worobey, et al. 2014a) | DNA | Influenzavirus A | 351 | 2280 | 3 |
| 66 | Wu_2018a | (Wu, et al. 2018) | (Wu, et al. 2019) | AA | mammalia | 90 | 3050199 | 5162 |
| 67 | Wu_2018b | (Wu, et al. 2018) | (Wu, et al. 2019) | DNA | mammalia | 90 | 9150597 | 15486 |

#### Table A.2. The probability distributions and their probability density function that we used for the Kolmogorov-Smirnov test.

| distribution | PDF | notes |
| --- | --- | --- |
| Alpha | $\frac{1}{x^{2}\Phi(\alpha)\sqrt{2\pi}}e^{-\frac{1}{2}{(\alpha-\frac{1}{x})}^{2}}$ | $\Phi$ is the normal CDF  α is the shape parameter |
| Beta | $\frac{{\Gamma(\alpha+\beta)x}^{\alpha-1}{(1-x)}^{\beta-1}}{\Gamma(\alpha)\Gamma(\beta)}$ | α and β are the shape parameters Γ is the gamma function |
| Bradford | $\frac{\alpha}{log(1+\alpha)(1+\alpha x)}$ | α is the shape parameter |
| Chi | $\frac{1}{2^{\alpha/2-1}\Gamma(\frac{\alpha}{2})}{x^{\alpha-1}e}^{-\frac{x^{2}}{2}}$ | α is the degrees of freedom  Γ is the gamma function |
| Chi-squared | $\frac{1}{2^{\alpha/2}\Gamma(\frac{\alpha}{2})}{x^{\alpha/2-1}e}^{-\frac{x}{2}}$ | α is the degrees of freedom  Γ is the gamma function |
| Double gamma | $\frac{\alpha}{2\Gamma(\alpha)}{\vert x\vert}^{\alpha-1}e^{-\vert x\vert}$ | α is the shape parameter  Γ is the gamma function |
| Double Weibull | ${\frac{\alpha}{2}\vert x\vert}^{\alpha-1}e^{-\left\vert x \right\vert^{\alpha}}$ | α is the shape parameter |
| Exponential normal | $\frac{1}{2\alpha}e^{\frac{1}{2\alpha^{2}}-\frac{x}{\alpha}}erfc(-\frac{x-\frac{1}{\alpha}}{\sqrt{2}})$ | α is the shape parameter |
| Exponential Weibull | ${\alpha\beta(1-e^{{-x}^{\beta}})}^{\alpha-1}e^{{-x}^{\beta}}x^{\beta-1}$ | α and β are the shape parameters |
| Exponential power | $\alpha x^{\alpha-1}e^{1+x^{\alpha}-e^{x^{\alpha}}}$ | α is the shape parameter |
| Gamma |  |  |
| Generalized logistic | $\alpha\frac{e^{-x}}{{(1+e^{-x})}^{\alpha+1}}$ | α is the shape parameter |
| Generalized Pareto | ${(1+\alpha x)}^{-1-\frac{1}{\alpha}}$ | α is the shape parameter |
| Generalized normal | $\frac{\alpha}{2\Gamma(1/\alpha)}e^{-\left\vert x \right\vert^{\alpha}}$ | α is the shape parameter  Γ is the gamma function |
| Generalized exponential | $(\alpha+\beta(1-e^{\gamma x}){)e}^{-\alpha x-\beta x+\frac{\beta}{\gamma}(1-e^{-\gamma x})}$ | α, β, γ are the shape parameters |
| Generalized gamma | $\frac{{\vert\beta\vert x}^{\alpha\beta-1}}{\Gamma(\alpha)}e^{-x^{\beta}}$ | α and β are the shape parameters |
| Half-logistic | $\frac{{2e}^{-x}}{{(1+e^{-x})}^{2}}$ |  |
| Half-normal | $\sqrt{2/\pi}e^{\frac{-x^{2}}{2}}$ |  |
| Upped half of the generalized normal | $\frac{\alpha}{\Gamma(1/\alpha)}e^{-\left\vert x \right\vert^{\alpha}}$ | α is the shape parameter |
| Inverse-gamma | $\frac{x^{-\alpha-1}}{\Gamma(\alpha)}e^{-\frac{1}{x}}$ | α is the shape parameter |
| Inverse-normal | $\frac{1}{\sqrt{2\pi x^{3}}}e^{-\frac{{(x-\alpha)}^{2}}{2x\alpha^{2}}}$ | α is the shape parameter |
| Inverse-Weibull | ${\alpha x}^{-\alpha-1}e^{-x^{-\alpha}}$ | α is the shape parameter |
| Laplace | $\frac{1}{2}e^{-\vert x\vert}$ |  |
| Log-gamma | $\frac{e^{\alpha x-e^{x}}}{\Gamma(\alpha)}$ | α is the shape parameter  Γ is the gamma function |
| Logistic | $\frac{e^{-x}}{{(1+e^{-x})}^{2}}$ |  |
| Log-Laplace | $\frac{\alpha}{2}x^{\alpha-1} 0<x<1$  $\frac{\alpha}{2}x^{-\alpha-1} x\geq1$ | α is the shape parameter |
| Log-normal | $\frac{1}{\alpha x\sqrt{2\pi}}e^{-\frac{({\log x)}^{2}}{2\alpha^{2}}}$ | α is the shape parameter |
| Maxwell | $\sqrt{2/\pi}{x^{2}e}^{\frac{-x^{2}}{2}}$ |  |
| Normal | $\frac{1}{\sqrt{2\pi}}e^{-\frac{x^{2}}{2}}$ |  |
| Pareto | $\frac{\alpha}{x^{\alpha+1}}$ | α is the shape parameter |
| Power-law | ${\alpha x}^{\alpha-1}$ | α is the shape parameter |
| Power Log-normal | $\frac{\alpha}{x\beta}\phi(\frac{\log x}{\beta}){(\Phi\left( -\frac{\log x}{\beta} \right))}^{\alpha-1}$ | α and β are the shape parameters  $\phi$ is the normal PDF  $\Phi$ is the normal CDF |
| Power normal | $\alpha\phi(x){(\Phi\left( -x \right))}^{\alpha-1}$ | α is the shape parameter  $\phi$ is the normal PDF  $\Phi$ is the normal CDF |
| Uniform | 1 |  |
| Weibull maximum | ${\alpha(-x)}^{\alpha-1}e^{-{(-x)}^{\alpha}}$ | α is the shape parameter |
| Weibull minimum | ${\alpha x}^{\alpha-1}e^{-x^{\alpha}}$ | α is the shape parameter |

Gamma function: $\Gamma\left( \alpha\right)= \int_{0}^{\infty} x^{\alpha-1}e^{-x}dx=\left( \alpha-1 \right)!$

### Figures

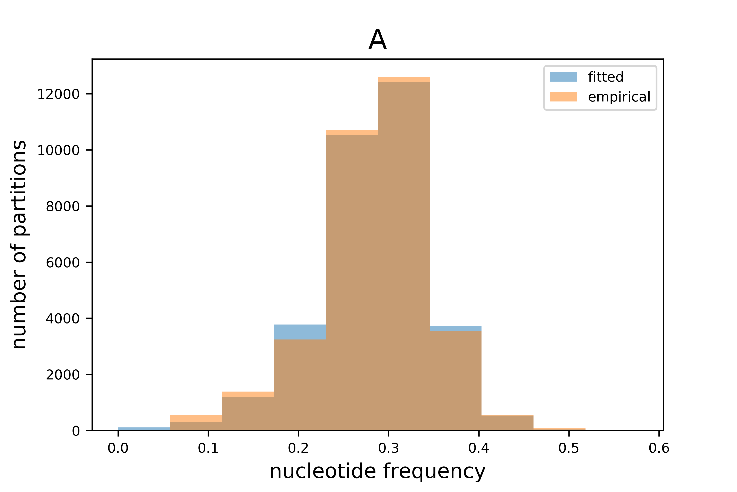

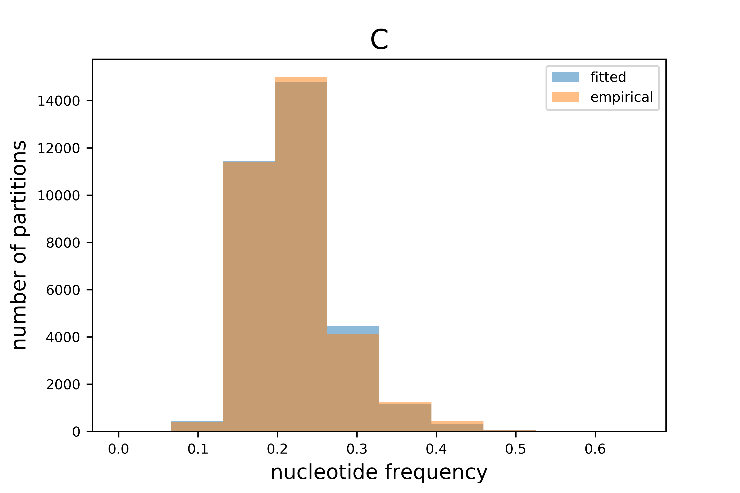

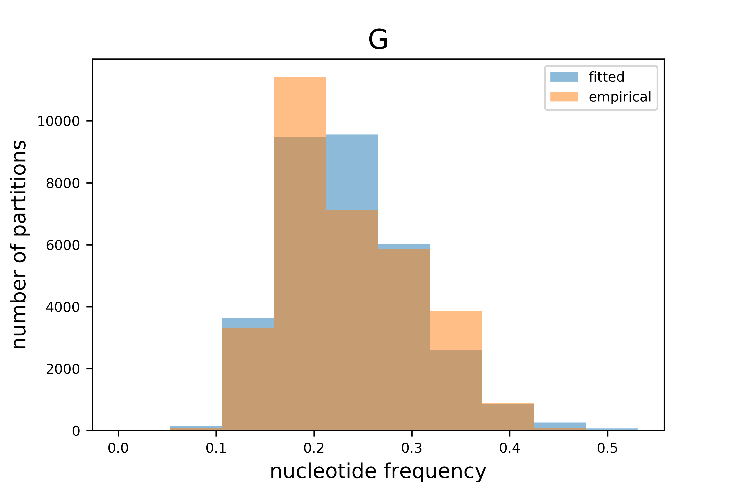

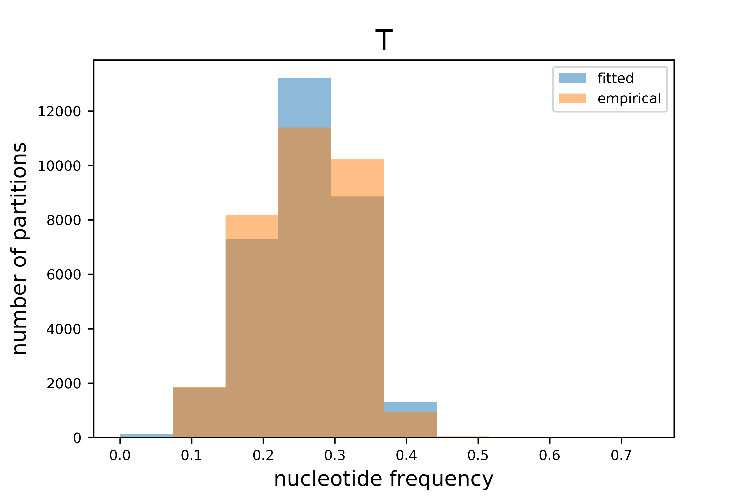
Figure A.1.	distribution of the nucleotide frequencies. The empirical nucleotide frequencies for each single partition were estimated using IQ-TREE (orange) and the Fitted distribution (blue) were sampled from the best-fit distribution with the same number of partitions.

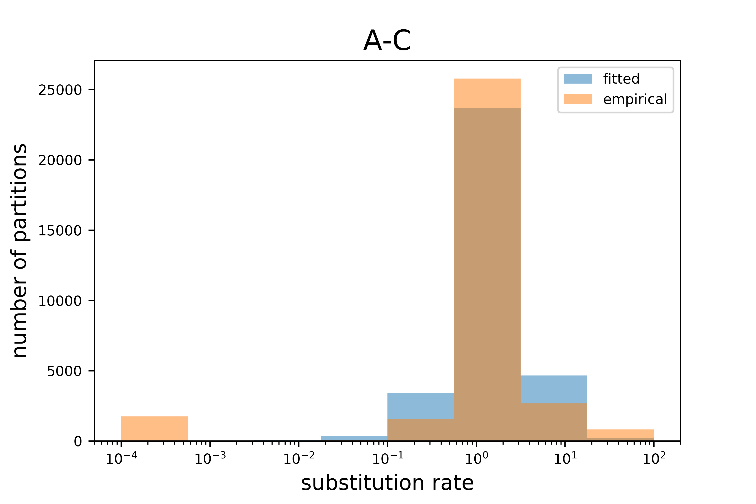

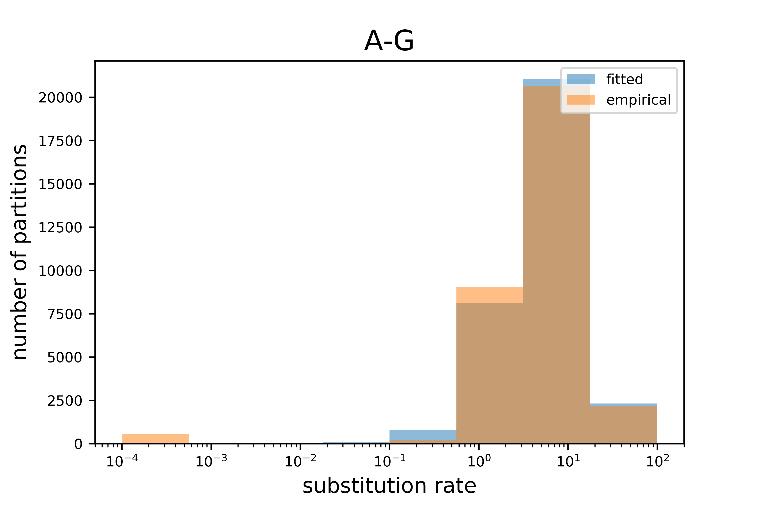

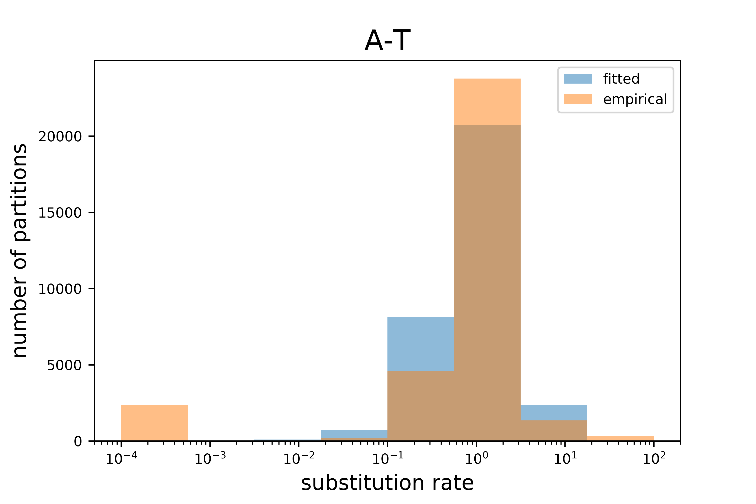

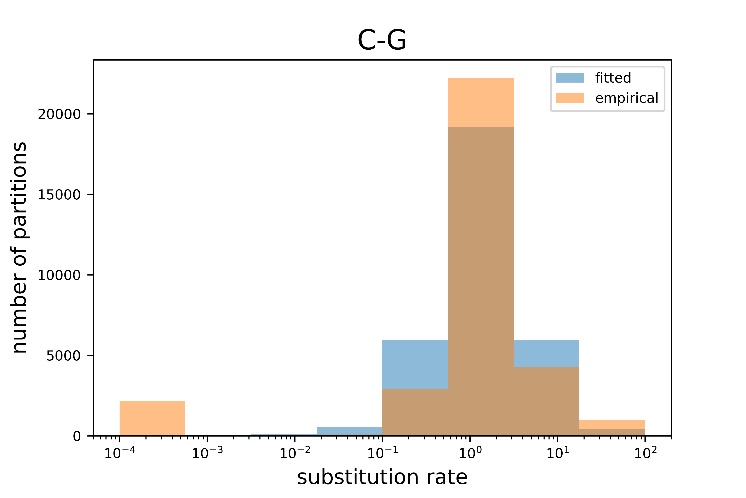

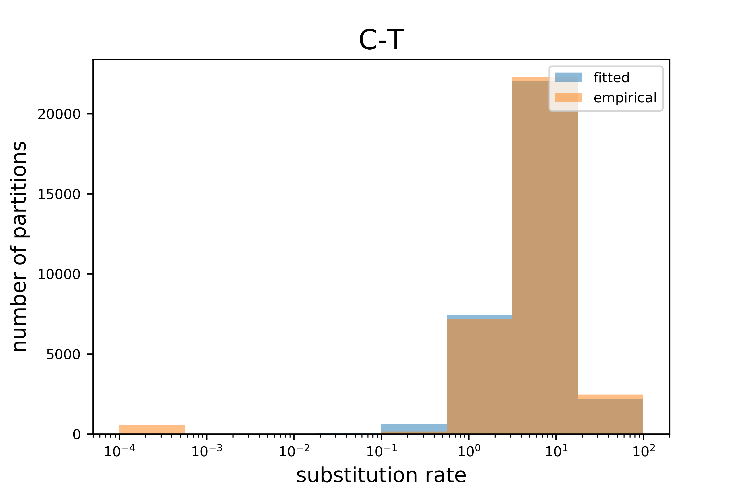

#### Figure A.2. distribution of the GTR parameters. The best-fit substitution rate matrix for each single partition was estimated using IQ-TREE (orange) and the Fitted distribution (blue) were sampled from the best-fit distribution with the same number of partitions.

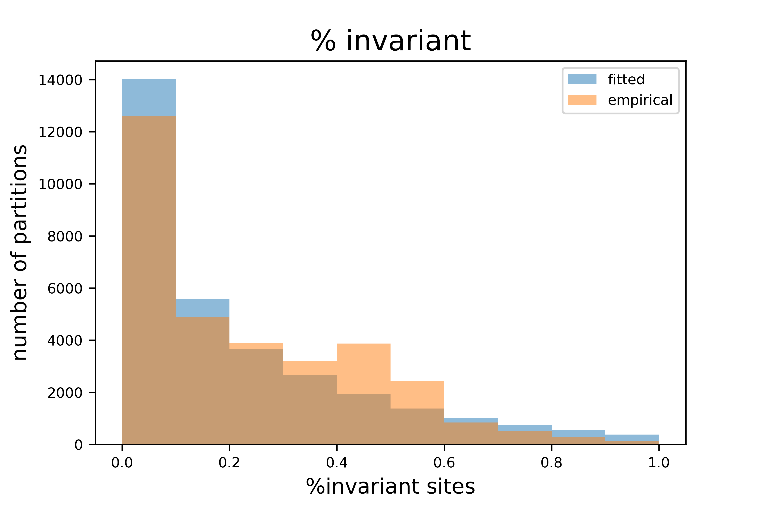

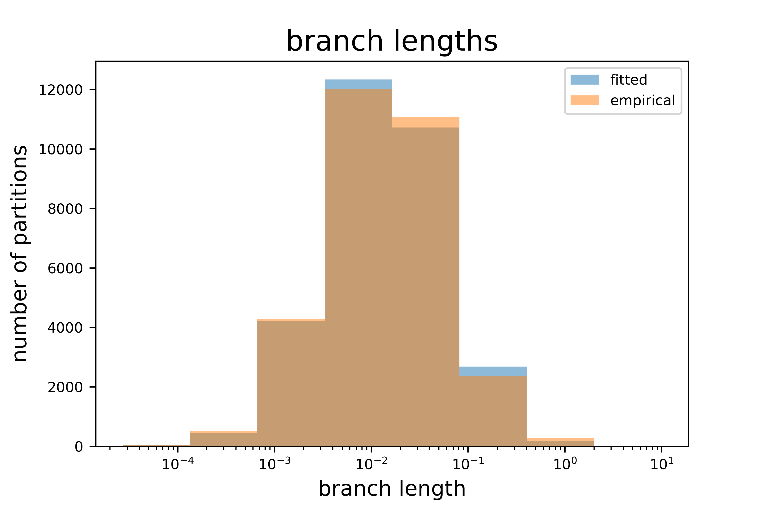

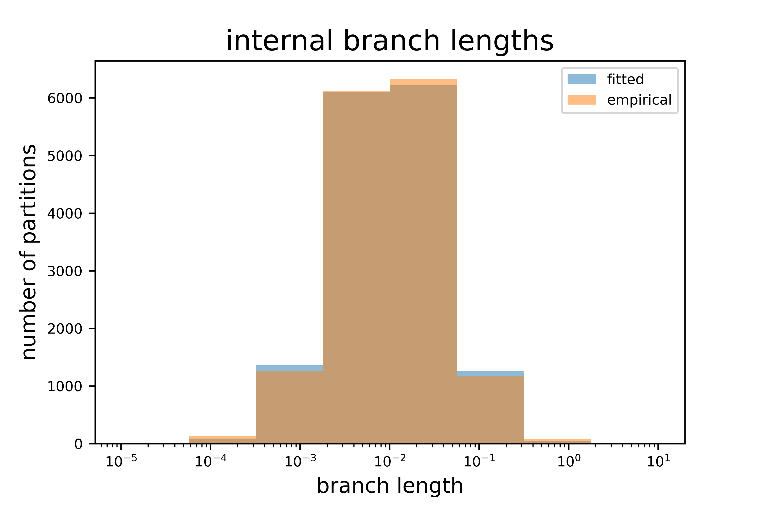

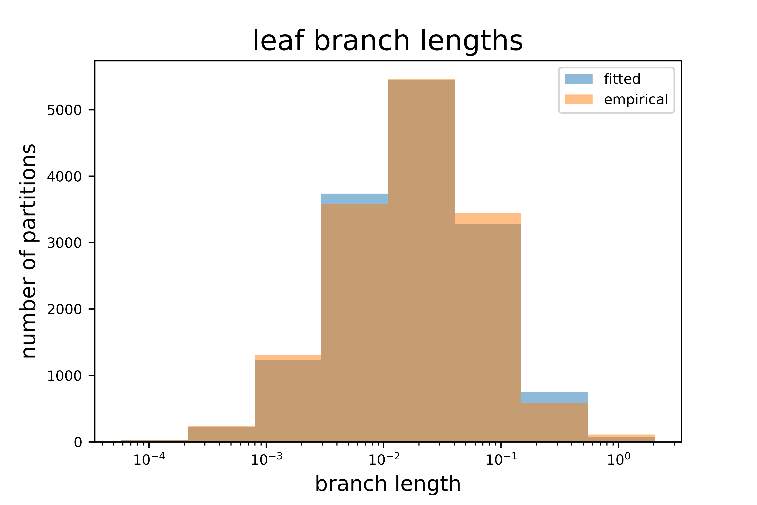

#### Figure A.3. The distribution of branch lengths and proportion of invariant sites

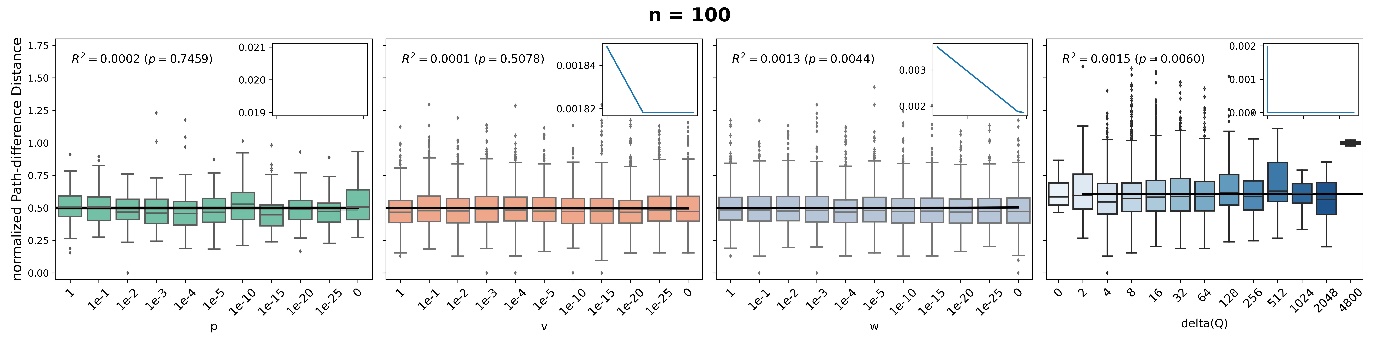

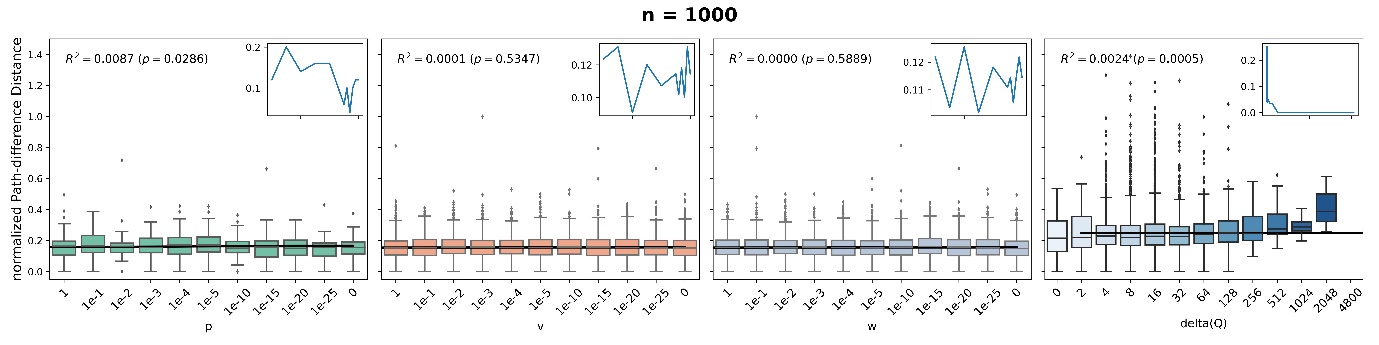

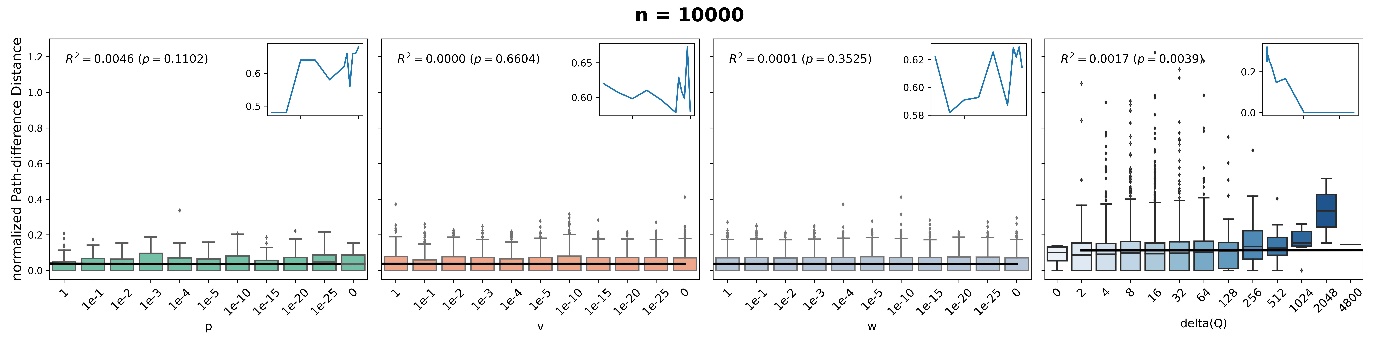

(a)

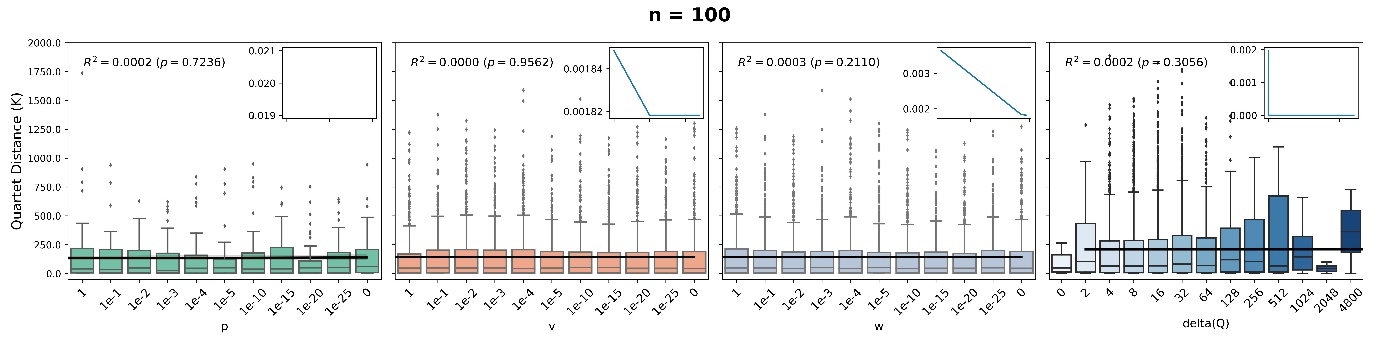

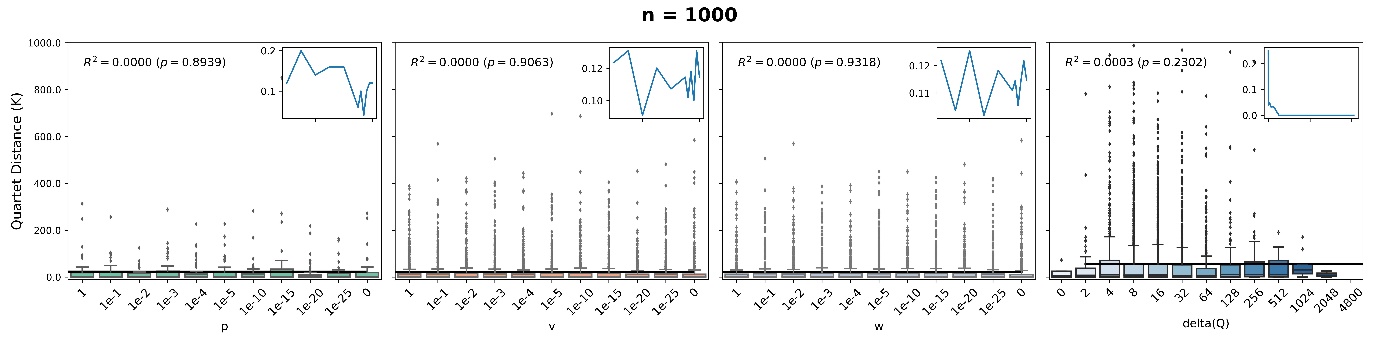

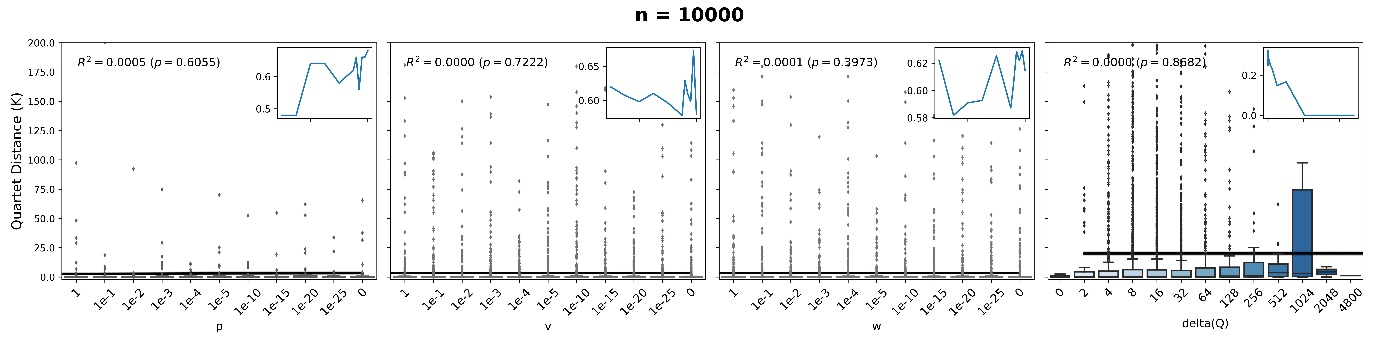

(b)

#### Figure A.4. (a) Normalized Path-difference and (b) Quartet metrics between the estimated tree topology and the original tree topology as a function of the inheritance weight (ν, ω ,ρ) and the distance between the two matrices. The small plots show the proportion of datasets in which the distance between the estimated topology and the original topology equals to zero as a function of the inheritance weight and the distance between the two matrices.

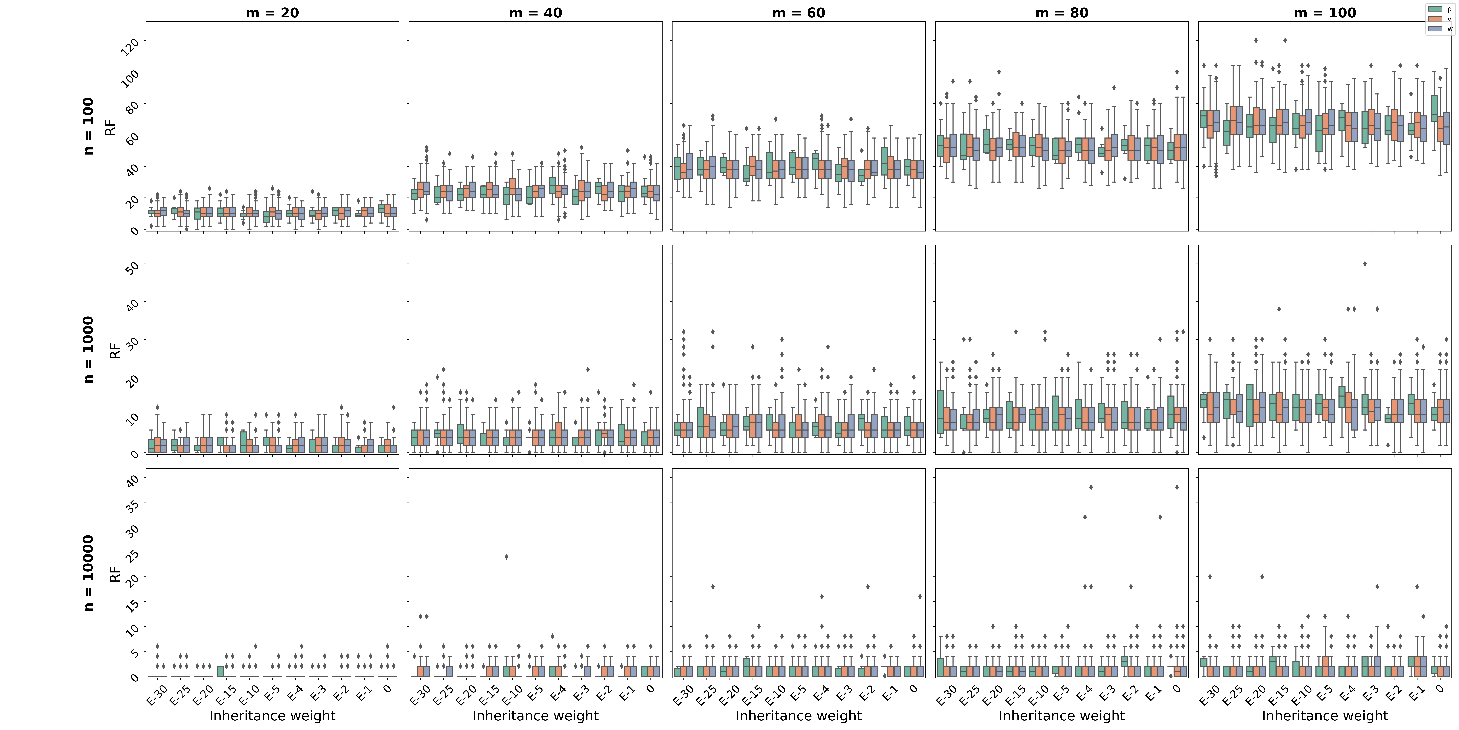

#### Figure A.5. The Robinson-Foulds metric as a function of the inheritance weight for each number of taxa (m) and number of site (n).

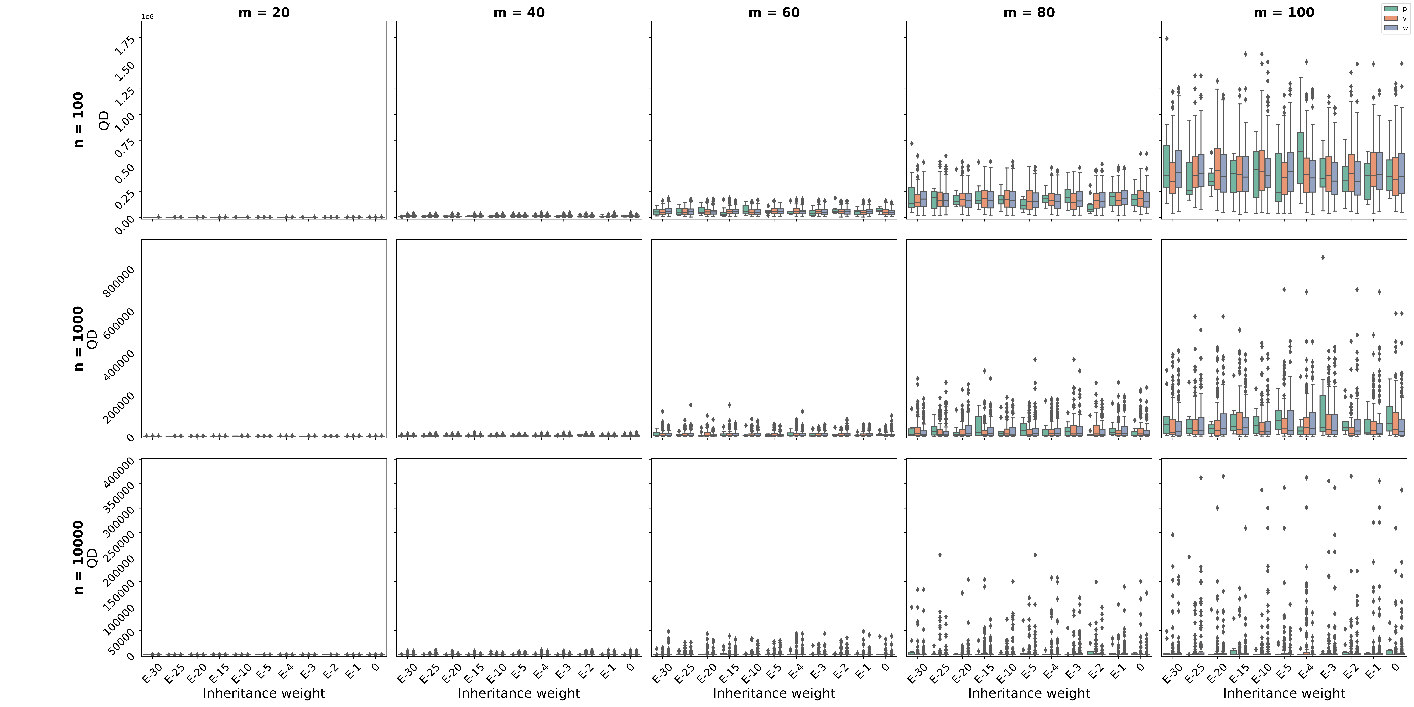

#### Figure A.6. The Quartet distance as a function of the inheritance weight for each number of taxa (m) and number of site (n).

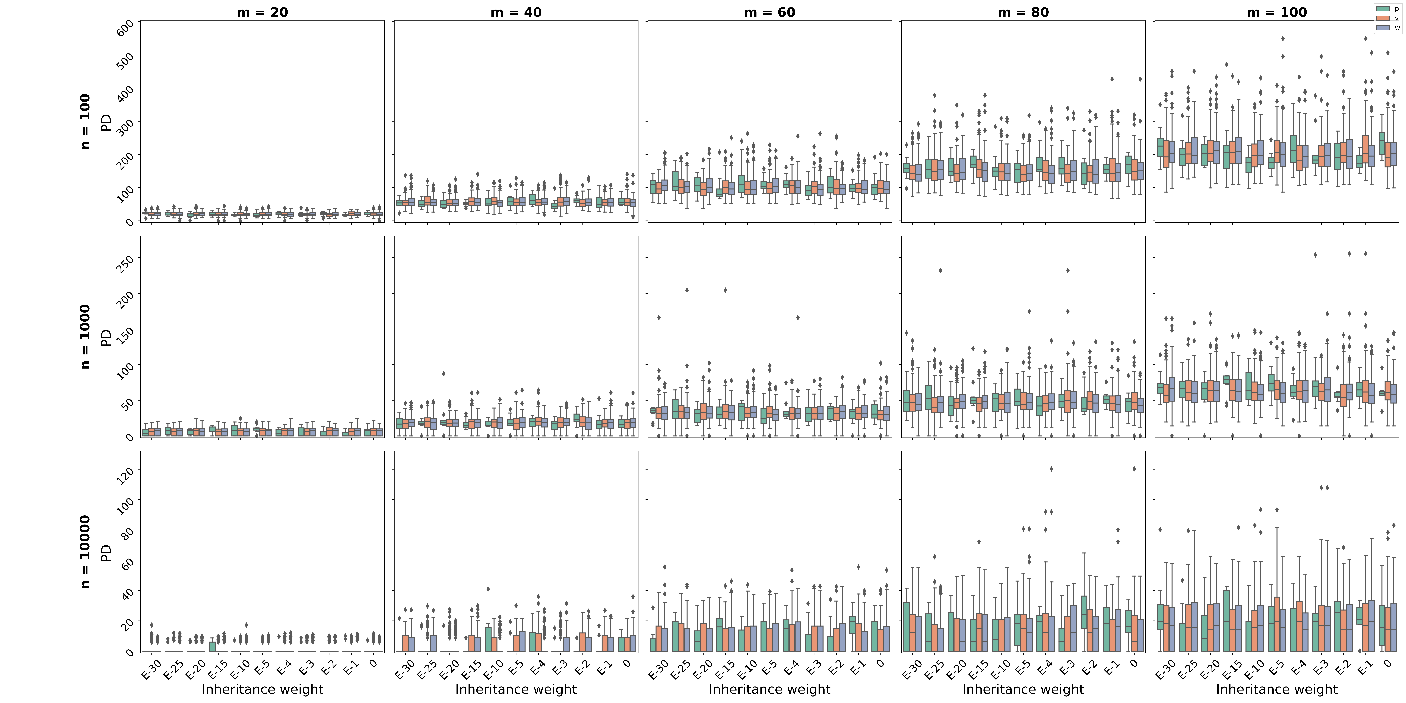

#### Figure A.7. The Path-Difference distance as a function of the inheritance weight for each number of taxa (m) and number of site (n).

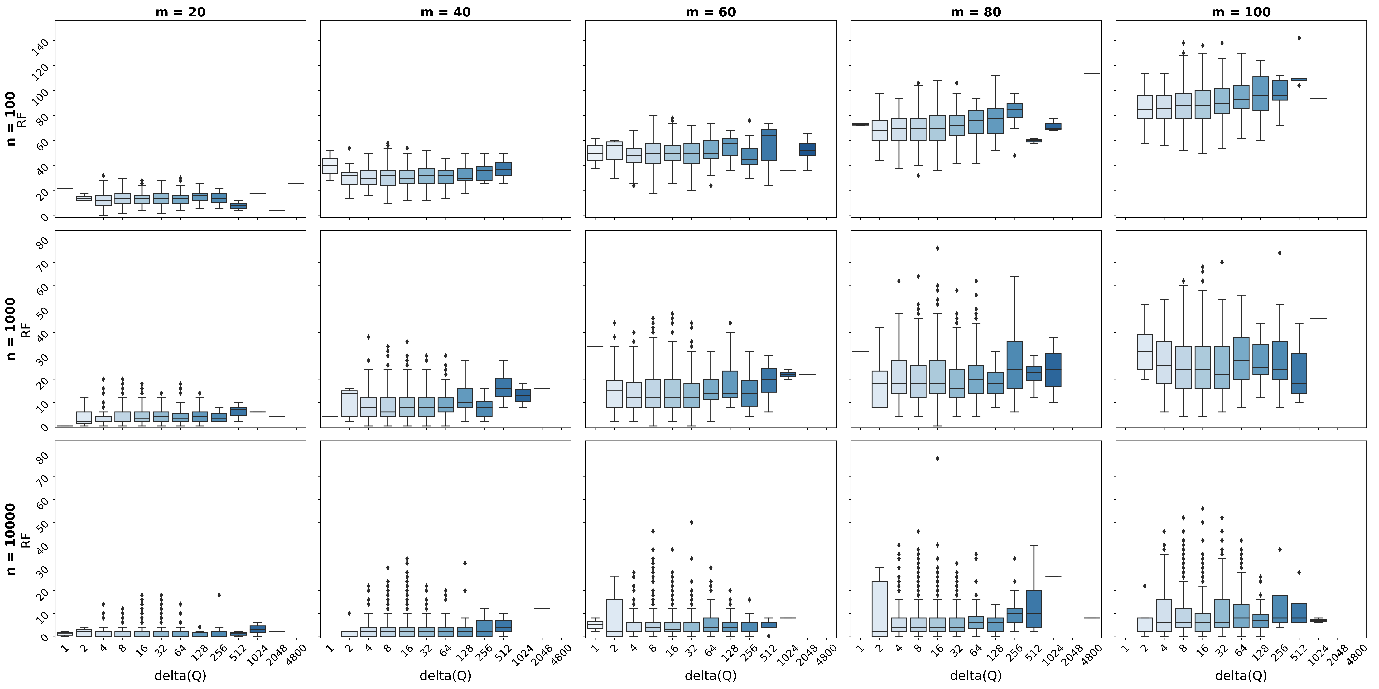

#### Figure A.8. The Robinson-Foulds metric as a function of the maximum Euclidian distance between the two matrices for each number of taxa (m) and number of site (n).

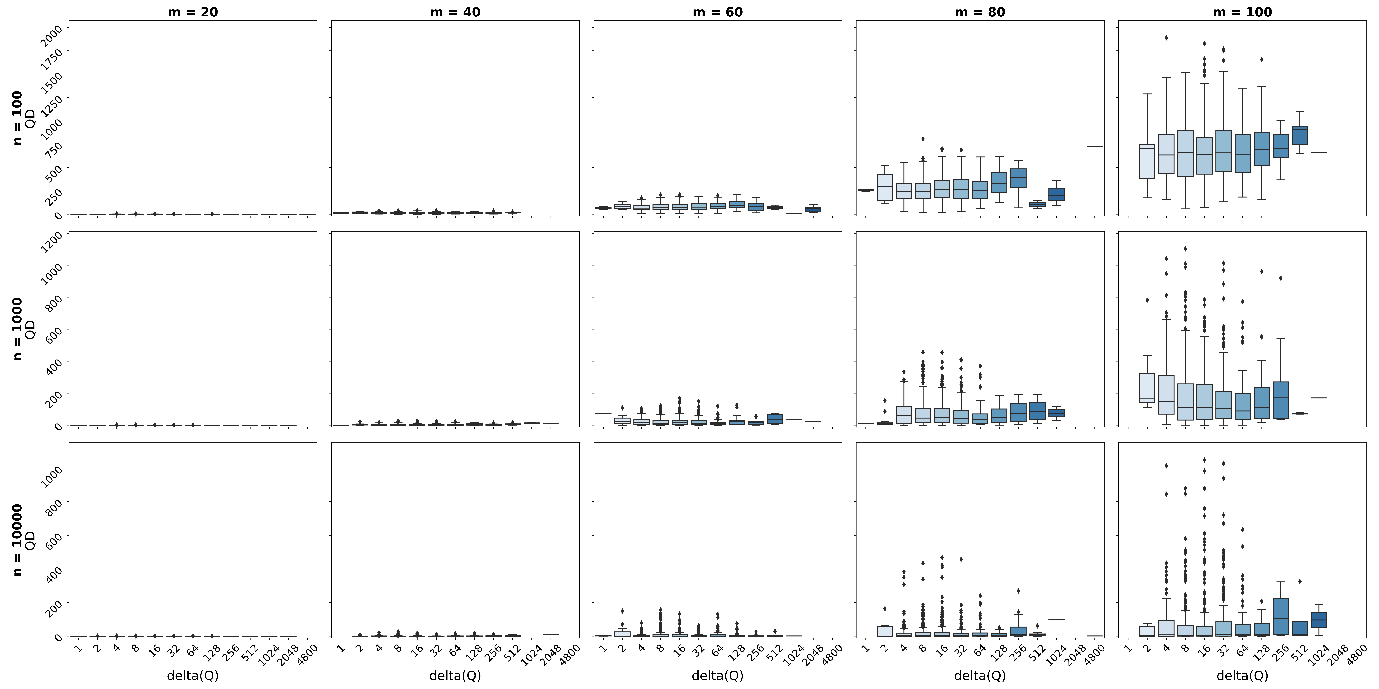

#### Figure A.9. The Quartet distance as a function of the maximum Euclidian distance between the two matrices for each number of taxa (m) and number of site (n).

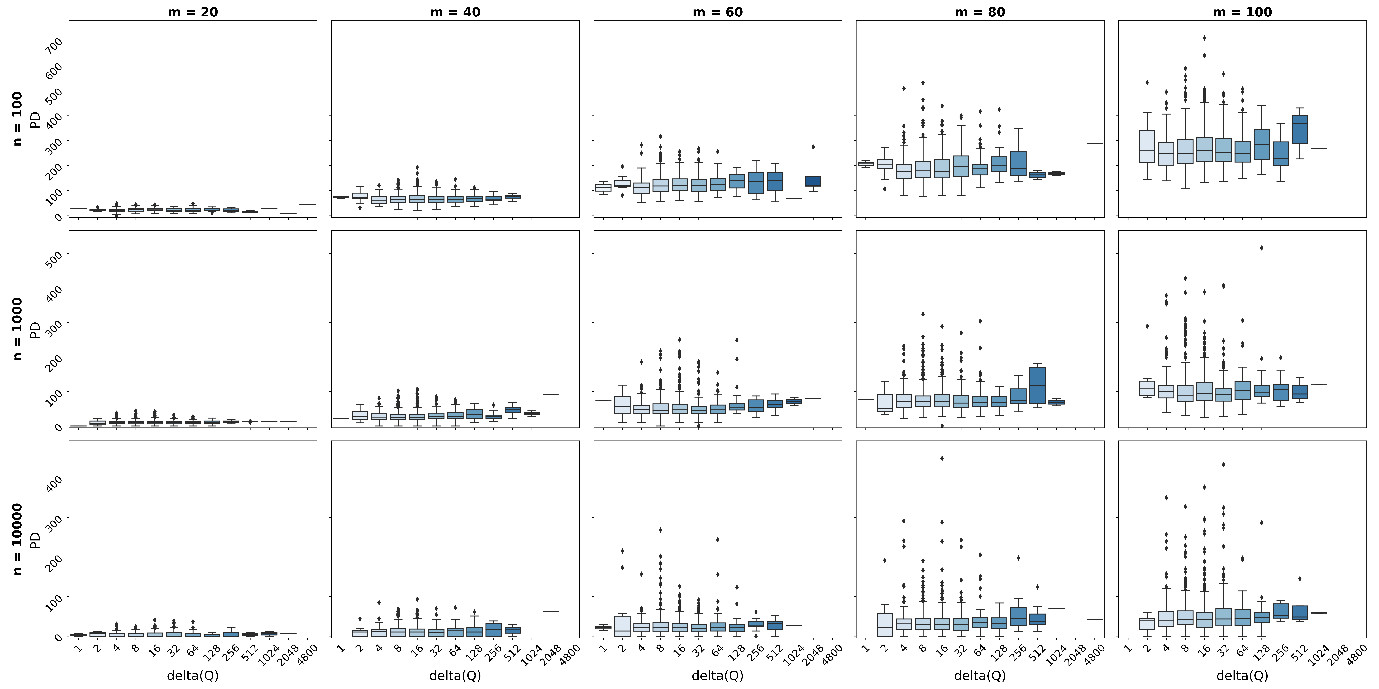

#### Figure A.10. The Path-Difference distance as a function of the maximum Euclidian distance between the two matrices for each number of taxa (m) and number of site (n).

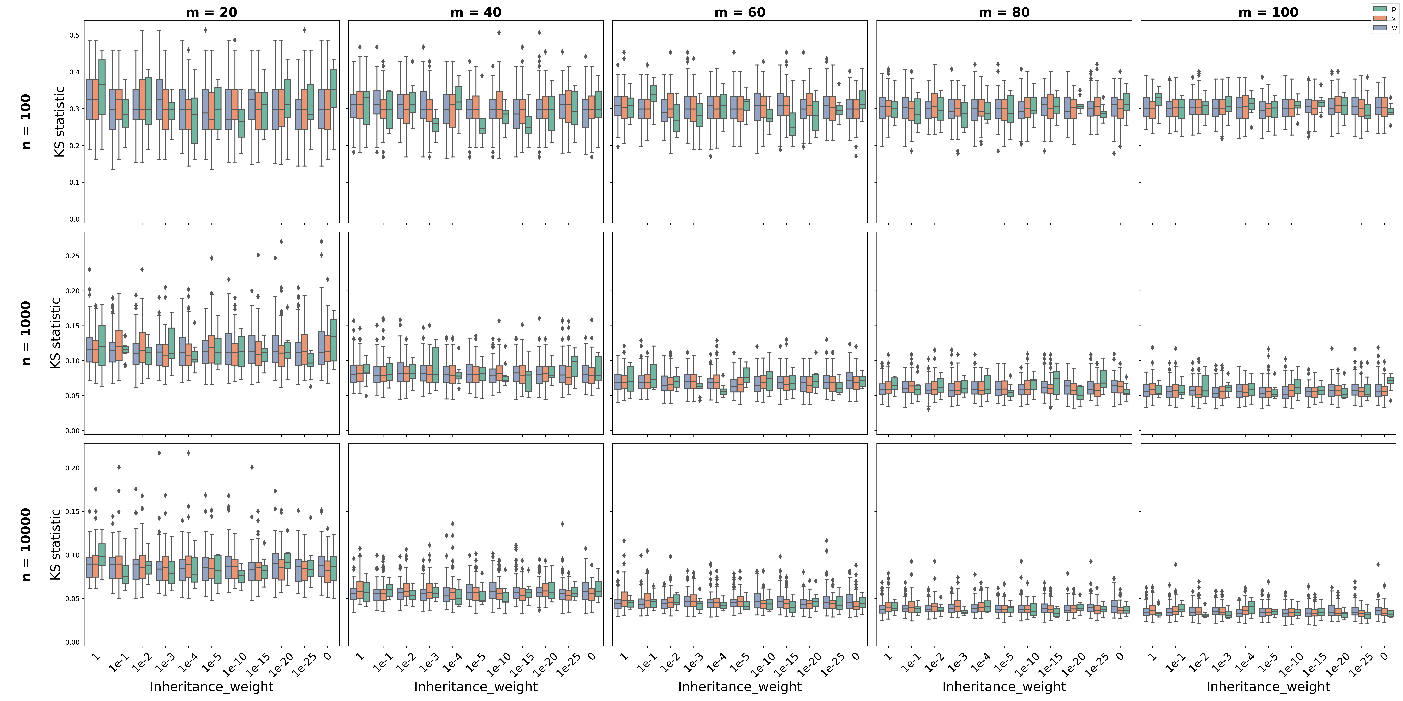

#### Figure A.11. Two-sample Kolmogorov-Smirnov test statistic as a function of the inheritance weight of the base frequencies (ν), the substitution model (ρ), the inheritance weight of the substitution rates (ω) for each number of taxa (m) and number of site (n).

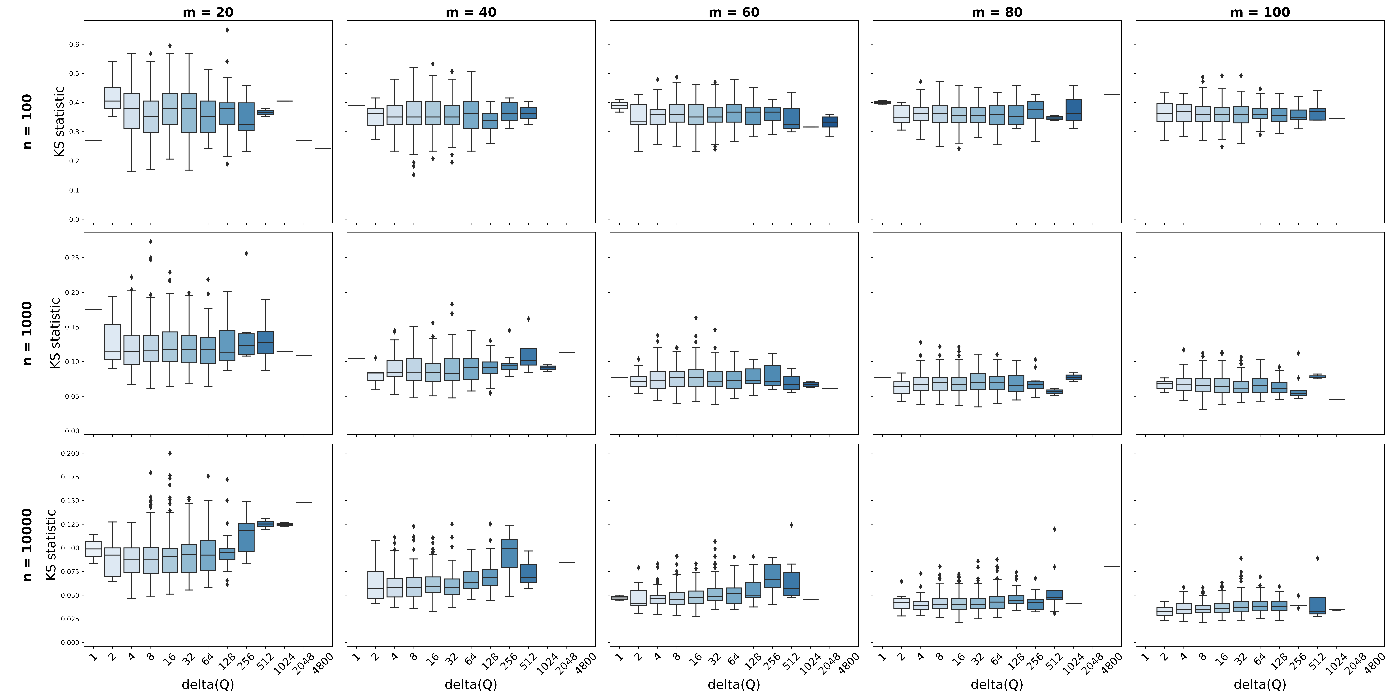

#### Figure A.12. Two-sample Kolmogorov-Smirnov test statistic as a function of the inheritance weight of the Euclidian distance between the two matrices for each number of taxa (m) and number of site (n).
